# Tau depletion in human neurons mitigates Aβ-driven toxicity

**DOI:** 10.1101/2022.12.23.521772

**Authors:** Bryan Ng, Jane Vowles, Dayne Beccano-Kelly, M Irina Stefana, Darragh P. O’Brien, Nora Bengoa-Vergniory, Feodora Betherat, Ajantha Abey, Philippa Carling, Peter Kilfeather, John A. Todd, Tara M. Caffrey, Natalie Connor-Robson, Sally A. Cowley, Richard Wade-Martins

## Abstract

**Background:** Alzheimer’s disease (AD) is an age-related neurodegenerative condition and the most common type of dementia, characterised by pathological accumulation of extracellular plaques and intracellular neurofibrillary tangles that mainly consist of amyloid-β (Aβ) and hyperphosphorylated tau aggregates, respectively. Previous studies in mouse models with a targeted knock-out of the microtubule-associated protein tau *(Mapt)* gene demonstrated that Aβ-driven toxicity is tau-dependent. However, human cellular models with chronic tau lowering remain unstudied.

**Methods:** In this study, we generated stable tau-depleted human induced pluripotent stem cell (iPSC) isogenic panels from two healthy individuals using CRISPR-Cas9 technology. We then differentiated these iPSCs into cortical neurons *in vitro* in co-culture with primary rat cortical astrocytes before conducting electrophysiological and imaging experiments for a wide range of disease-relevant phenotypes. Both AD brain derived and recombinant Aβ were used in this study to elicit toxic responses from the iPSC-derived cortical neurons.

**Results:** We showed that tau depletion in human iPSC-derived cortical neurons caused considerable reductions in neuronal activity without affecting synaptic density. We also observed neurite outgrowth impairments in two of the tau-depleted lines used. We found axonal transport of mitochondria, mitochondrial function, and cortical neuron differentiation propensity remained unaffected regardless of tau expression levels. Finally, tau depletion protected neurons from adverse effects mitigating the impact of exogenous Aβ-induced hyperactivity, deficits in retrograde axonal transport of mitochondria, and neurodegeneration.

**Conclusions:** Our study established stable human iPSC isogenic panels with chronic tau depletion from two healthy individuals. Cortical neurons derived from these iPSC lines showed that tau is essential in Aβ-driven hyperactivity, axonal transport deficits, and neurodegeneration, consistent with studies conducted in *Mapt-/-* mouse models. These findings highlight the protective effects of chronic tau lowering strategies in AD pathogenesis and reinforce the potential in clinical settings. The tau-depleted human iPSC models can now be applied at scale to investigate the involvement of tau in disease-relevant pathways and cell types.

## Background

Alzheimer’s disease (AD) is the most common age-related neurodegenerative disease and cause of dementia, and is characterised by pathological accumulations of two key proteins – amyloid-β (Aβ) and tau – in the brain (1). Accumulation of tau and Aβ remains the key pathology specific to AD as set out by the latest biological definition of AD in research from a National Institute on Aging-Alzheimer’s Association working group (2), although recent advances in research have identified additional systems, cell types and molecular changes that are involved in AD pathogenesis. However, the precise link between Aβ, tau and eventual neurodegeneration remains only partially understood.

Over the past few decades, researchers have shown in longitudinal studies that Aβ accumulation precedes tau pathology in AD brains and that AD brain-derived soluble Aβ induces tau hyperphosphorylation which precedes tau aggregation (3, 4). Furthermore, disease-causing mutations in familial AD are associated with proteins involved in the pathway of Aβ production i.e., amyloid precursor protein, presenilin 1 and presenilin 2, leading to the “amyloid cascade hypothesis” (5, 6). To investigate how tau fits into this hypothesis, mouse models with a targeted knock-out of the microtubule-associated protein tau *(Mapt)* gene have been used to demonstrate that Aβ-driven toxicity is tau-dependent as reported in studies examining neuronal activity, axonal transport, and neurodegeneration, amongst others (7–10). *Mapt-/-* mice are viable and do not present overt cognitive or behavioural deficits, although certain strains were reported to suffer from neurite outgrowth impairment and age-dependent motor dysfunction (10–12).

Even though mouse models have been integral in elucidating mechanisms of AD pathogenesis, distinct biological differences remain between human and mouse in the context of AD. Aged mice do not develop AD pathology without the introduction of disease-causing mutations indicating a different disease susceptibility as compared to humans (14). Even with the introduction of mutations found in familial AD cases, AD mouse models do not present with tau pathology as would be the case for typical AD pathological manifestation. One important difference between human and mouse biology is that only four-repeat tau isoforms, representing approximately half of all tau isoforms in a human brain, are expressed in adult mice (15). It is therefore imperative to assess the role of tau in the context of Aβ-driven toxicity in a human neuronal model in addition to studies involving mouse models. However, human *MAPT-/-* cell lines have not been studied in detail to better understand tau-dependent AD pathophysiology in a human genetic background.

Induced pluripotent stem cells (iPSCs) have emerged as a versatile human cell model which can be differentiated *in vitro* into almost any cell type, including neurons which are normally inaccessible without invasive surgeries (16). iPSCs can be genetically modified to represent a genotype of interest *in vitro*. Here, we generated stable *MAPT-/-* iPSC lines from two healthy individuals by clustered regularly interspaced short palindromic repeats (CRISPR)-Cas9-mediated gene editing. These iPSC lines were differentiated into cortical neurons and were examined for the effects of tau depletion in phenotypes that included neuronal activity, synapse loss, axonal transport, neurite outgrowth and neurodegeneration. We found that tau depletion caused significant reduction in neuronal activity but protected iPSC-derived cortical neurons from Aβ- driven neuronal hyperactivity, axonal transport deficits, and neurodegeneration.

## Methods and materials

All reagents were purchased from Sigma-Aldrich (Merck), and all cell cultures were maintained at 37°C with 5% CO2 in a humidified incubator unless stated otherwise.

### Generation MAPT-/- iPSC lines using CRISPR-Cas9

Two strategies targeting either Exon 1 or Exon 4 of the *MAPT* gene were employed in two separate previously published iPSC lines derived from dermal fibroblasts from healthy individuals: SBAd-03-01 (Exon 1 targeted; 31 years old female (17); *APOE ε2/ε3*); and SFC856-03-04 (Exon 4 targeted; 76 years old female (18); *APOE ε3/ε4*). The genetic manipulations of *MAPT* gene were achieved by using an Alt-R CRISPR- Cas9 System (Integrated DNA Technologies) following manufacturer’s protocol. The Cas9 Nuclease V3 was used with a single gRNA to result in a double-stranded break in Exon 1, whereas Exon 4 was targeted with a pair of gRNAs to create a 25 bp excision (Supplementary Table 1). To deliver hybridised Cas9-gRNA ribonucleoprotein into the iPSCs, a Neon Transfection System (Thermo) was used to transfect a suspension of 220,000 iPSCs with a single pulse at 1400 V with 20 ms pulse width. The transfected iPSCs were then plated onto one well of a 24-well plate for recovery until the cells became confluent. Thereafter, the cells were singularised before plating 2,000 cells per well in a 6-well plate on irradiated mouse embryonic fibroblast feeder cell line CF1. Ninety-six individual iPSC colonies were subsequently picked and transferred separately into a 96-well plate before they achieved confluence and were lysed for DNA extraction.

### Identification of two panels of isogenic MAPT-/- iPSC lines

Genomic DNA was extracted from each iPSC colony using a commercial kit (Qiagen), before the DNA samples were amplified using the BigDye^TM^ v3.1 reagent and sequenced with primers targeting either Exon 1 or Exon 4 spanning the gRNA binding sites (Supplementary Table 1). Before sending the DNA samples (10 μL per iPSC clone) for sequencing, samples were precipitated by adding 2 μL of 125 mM EDTA, 2 μL of 3 M sodium acetate and 50 μL of 100% ethanol to each sample. The samples were briefly vortexed before centrifugation at 3,000 g at 4°C for 30 min. After removing the supernatant by centrifugation at 185 g for 15 s, the DNA precipitates were washed with 70 μL of 70% ethanol per sample.

The sequencing results were analysed with the SnapGene^®^ Viewer v5.2.4 and BLAST^®^ (19) to identify candidate iPSC clones that underwent successful genetic modifications at the targeted regions. These candidate iPSC clones were further sub- cloned into pGEM^®^-T Easy Vectors (Promega) to determine the genetic modifications at the allelic level using the same primers for the Exon 1 and 4 targeted regions. The vectors were then transformed into One Shot^TM^ TOP10 Chemically Competent *E. coli* (Thermo) by heat shock at 42°C for 45 s. At least six bacterial clones were picked and sequenced the way it was described above using T7 and SP6 promoter primers (embedded in pGEM^®^-T Easy Vectors).

Once the homozygosity/heterozygosity of the intended genetic modifications was determined, the sequence-confirmed iPSC clones were differentiated into neural progenitor cells (NPCs) or neurons and their cell lysates were analysed by western blot to confirm tau depletion. A Human OmniExpress v1.2 BeadChip array (Illumina) was used post-editing to check for any gross karyotype abnormalities, and none were detected in all iPSC lines used in this study (Supplementary Fig. 1).

### Lentivirus generation

The lentiviral particles used in this study expressing genes of interest were generated by expressing lentiviral proteins in HEK293T cells grown in DMEM/F12 medium supplemented with 10% FBS, 2 mM L-glutamine, 100 μM non-essential amino acids and 50 U/ml Penicillin-Streptomycin (all from Thermo). HEK293T cells were passaged with TrypLE^TM^ (Thermo) and seeded at 80,000 cells/cm^2^ (1,000,000 cells in a T75 flask) 24 h before plasmid transfection.

The following protocol describes the reagents used for a single T75 flask of HEK293T cells. A Lipofectamine^TM^ 3000 transfection reagent (Thermo) was used to deliver plasmids into HEK293T cells. 40 μL Lipofectamine^TM^ 3000 transfection reagent was first mixed with 50 μL of P3000 reagent (from the same kit) in 1 mL of Opti-MEM^TM^ medium (Thermo) for at least 5 min. Second-generation lentiviral packaging plasmids psPAX2 (#12260; 10 μg) and pMD2.G (#12259; 5.65 μg) were mixed with plasmids expressing genes of interest (FUdeltaGW-rtTA (rtTA; #19780; 9.6 μg), pTet-O-Ngn2- puro (Ngn2; #52047; 9.6 μg) or pLV-TetO-hNGN2-eGFP-Puro (Ngn2-GFP; #79823; 9.6 μg)) to achieve equimolar ratios in 1 mL of Opti-MEM^TM^ medium in another tube.

All plasmids were purchased from Addgene. Finally, the plasmid tube was added to the Lipofectamine tube dropwise before adding the mixture to the cells now grown in 13 mL of the above-mentioned medium without Penicillin-Streptomycin. The transfection reagents were left overnight and replaced with fresh medium the following day with Penicillin-Streptomycin.

The lentivirus-containing cell supernatant was collected 48 h after the last medium change, and it was centrifuged at 400 g for 5 min to remove any cells before filtering through a cellulose acetate membrane with 0.45 μm pore size (Millipore). The supernatant was snap-frozen in dry ice before storing it at -80°C. For the pLV-TetO- hNGN2-eGFP-Puro virus, the aliquots were thawed once and sub-aliquoted in small volume to keep the lentivirus titre consistent since a small amount of this lentivirus was needed per transduction.

### Differentiation of iPSC lines into cortical neurons

All iPSC lines were seeded on Matrigel^®^ (Corning)-coated plates and cultured in mTeSR^TM^ medium (STEMCELL Technologies) which was replaced daily. Fully confluent iPSC culture was passaged at a 1:6 ratio with TrypLE^TM^ (Thermo) for 5 min in the incubator and pelleted by centrifugation at 400 g for 5 min.

Two variations of cortical neuron differentiation protocols were adapted and combined from Shi *et al.* (pharmacologically directed) (20) and Zhang *et al.* (transcription factor directed) (21) into a single protocol and adapted as previously described (22). All media were filtered through a polyethersulfone membrane with a 0.22 μm pore size (Millipore). All cortical neuron cultures were seeded in co-culture with a confluent layer of primary rat cortical astrocytes (Thermo) except for harvesting cell lysate or live assays with dyes which are not cell-type specific. For the neuronal monoculture without rat astrocytes, the end point of differentiation was set to be Day 50 or before to ensure that the culture remained relatively homogeneous without any contamination with non-neuronal cells that might emerge sporadically throughout the differentiation process.

The first 25 days of cortical neuron differentiation was adapted from Shi et al. (20). All iPSC lines were first seeded concurrently on Geltrex^TM^ (Thermo)-coated plates at 105,000 cells/cm^2^ (1,000,000 cells in a well of a six-well plate) in mTeSR^TM^ medium supplemented with 10 μM Y-27632 (Tocris). The Y-27632 compound was removed from the medium the next day and subsequently the medium was replaced daily. When the iPSC culture became confluent (Day 0), it was washed once with phosphate-buffered saline (PBS) before changing to neural induction medium (NIM) which is neural maintenance medium (NMM) supplemented with 10 μM SB-431542 (Tocris) and 1 μM Dorsomorphin. NMM contains two types of media at 1:1 ratio: (1) Neurobasal^TM^ medium (Thermo) supplemented with 1X B-27^TM^ supplement (Thermo), 25 U/ml Penicillin-Streptomycin (Thermo) and 1 X GlutaMAX (Thermo) (2) DMEM/F12 medium (Thermo) supplemented with 1X N-2 supplement (Thermo), 50 μM non- essential amino acid (Thermo), 50 μM 2-mercaptoethanol (Thermo), 2.5 μg/ml insulin, 1 mM sodium pyruvate (Thermo) and 2 mM L-glutamine (Thermo). The NIM was subsequently replaced daily until Day 12.

The cells were passaged on Day 12 with 1 mg/ml dispase (10 mg/ml dispase dissolved in PBS and added directly to the cell culture media), for approximately 10 to 30 min in the incubator depending on the lifting efficiency. A sheet of cells would detach from the plate bottom and was then broken up by pipetting once before transferring to 10 mL of NIM. The cell aggregates were allowed to settle by gravity before they were washed by 10 mL of NIM twice. Finally, the cell aggregates were seeded onto 15 μg/ml fibronectin-coated plates at a 1:2 ratio in NIM.

On Day 13 and 15, the cells were provided with NMM supplemented with 20 ng/ml FGF2. On Day 17, the cell cultures were passaged as on Day 12 except that the cells were washed and plated in NMM. The cell aggregates were seeded onto 100 μg/ml poly-L-ornithine plus 10 μg/ml laminin (PO/L)-coated plates at a 1:3 ratio (the plates were first coated with PO overnight at 37°C, before they were washed once with PBS and then coated with laminin overnight). Finally, the NMM was replaced every two days until Day 25 where the NPCs were lifted by incubating with Accutase^®^ and cryo- preserved at 2,000,000 cells/vial in NMM supplemented with 20 ng/ml FGF2 and 10% dimethyl sulfoxide.

The second part of the differentiation protocol beyond Day 25 was modified from Zhang *et al.* (21) and the protocol relies on a doxycycline-inducible overexpression system of *Ngn2* to result in an excitatory cortical neuronal identity. Each vial of NPCs was thawed, spun down, and plated into a single well in a twelve-well plate (Day 25). *Ngn2* and *rtTA* lentiviruses (25 μL each) were added into the cell suspension in NMM (supplemented with 20 ng/ml FGF2 and 10 μM Y-27632) as the NPCs settled onto PO/L-coated plates. The medium was replaced 8 h later with fresh NMM. On Day 28, the NMM was replaced by neuronal medium consisting of Neurobasal^TM^ medium (Thermo) supplemented with 1X B27^TM^ (Thermo), 1X GlutaMAX^TM^ (Thermo), 25 U/ml Penicillin-Streptomycin (Thermo), 10 ng/ml neurotrophin-3, 10 ng/ml BDNF (Peprotech), 1 µg/ml doxycycline, 200 ng/ml laminin and 200 µM ascorbic acid. On Day 30, the neuronal medium was replaced and supplemented with 1 µg/ml puromycin.

On Day 32, the neurons were passaged with Accutase^®^ and seeded in co-culture with a confluent monolayer of primary rat cortical astrocyte seeded onto 10 µg/ml poly-D-lysine-coated plates (coated overnight and washed once with PBS before seeding) at its third passage (Thermo). The iPSC-derived neuron-rat astrocyte co-culture was subsequently half-fed with the neuronal medium only twice a week. On Day 40, the co-culture was treated with 100 nM Ara-C which was gradually removed over subsequent half-feedings. The neuronal culture was then allowed to age until Day 80 or after for downstream experiments.

When co-culture was not suitable for a particular experiment, the neurons were seeded onto PO/L-coated plates. The primary rat cortical astrocyte culture was maintained in DMEM with high glucose supplemented with 15% foetal bovine serum (FBS) (Thermo), 2 mM L-glutamine (Thermo) and 50 U/ml Penicillin-Streptomycin (Thermo). After the neurons were seeded onto the astrocyte, the co-culture was maintained in the neuronal medium only.

### Western blot

The NPCs were seeded on PO/L-coated plates on Day 25 and maintained until Day 30 for harvesting without virus transduction, while others were transduced with the *Ngn2* lentivirus and final-plated on Day 32 for harvesting on Day 50 as neurons. The cells were washed once with PBS before they were lysed on ice in radioimmunoprecipitation assay (RIPA) buffer containing 50 mM Tris-HCl, 150 mM NaCl, 1% (v/v) Triton X-100, 1% (w/v) sodium deoxycholate and 0.1% (w/v) sodium dodecyl sulphate (SDS) at pH 7.4. The RIPA buffer was supplemented with cOmplete^TM^ mini EDTA-free protease inhibitor cocktail (Roche). The lysate was then transferred into a tube on ice and left incubating for at least 15 min. Finally, the lysate was centrifuged at 20,000 g for 30 min at 4°C before the supernatant was frozen at - 80°C.

To quantify protein concentration in the lysates, a Pierce^TM^ Bicinchoninic Acid Protein Assay Kit (Thermo) was used following manufacturer’s instructions. Briefly, 12.5 μL of either bovine serum albumin (BSA) standards or cell lysates were added into a 96- well plate followed by 100 μL of the BCA reagent. The plate underwent incubation at 37°C for 30 min before it was read for absorbance at 562 nm wavelength in a PHERAstar^®^ microplate reader (BMG Labtech). Protein concentration was determined by comparing to a BSA standard curve.

For western blots to examine total tau protein, cell lysates underwent dephosphorylation with (final concentration in the mixture) 10% MnCl_2_, 10% reaction buffer and 0.38 μL lamda-phosphatase (New England Biolab; sufficient for up to 10 μg of cell lysate) for 1 h at 30°C prior to adding the sample loading dye for denaturation. Cell lysates were then denatured by heating at 95°C for 10 min with sample loading dye containing (final concentration in the mixture) 2% SDS, 5% 2-mercaptoethanol, 10% glycerol, 0.002% bromophenol blue and 375 mM Tris at pH 6.8.

Denatured samples were loaded into precast 4-15% gradient polyacrylamide gels (Bio-Rad) submerged in running buffer consisting of 25 mM Tris base, 192 mM glycine and 0.1% SDS. The gel was subject to 100 V for 3.5 h and the gel underwent semi-dry transfer by a Trans-Blot^®^ Turbo™ Transfer System (Bio-Rad) onto a 0.2 µm polyvinylidene fluoride membrane at 1.3 A for 11 min. The resultant blot was washed once in tris-buffered saline with 0.1% Tween-20 (TBST) and blocked in 5% milk dissolved in TBST shaking for 1 h at room temperature. The blot was then incubated with primary antibodies (Supplementary Table 2) diluted in 1% milk in TBST rocking overnight at 4°C.

The blot was washed with TBST shaking for 10 min at room temperature three times prior to incubation with secondary antibodies (Supplementary Table 2) diluted in 1% milk in TBST for 1 h at room temperature. Thereafter, the blot was washed for for three more times before adding a Immobilon^®^ Western Chemiluminescent HRP Substrate (Millipore) to the blot. The blot was imaged in the ChemiDoc^TM^ Imager (Bio-Rad) and analysed in the Image Lab v6.1 (Bio-Rad) software.

### Immunoprecipitation-mass spectrometry (IP-MS) analysis

All *MAPT+/+* and *MAPT-/-* iPSC lines from both isogenic panels were differentiated into Day 50 neurons but every line was maintained separately to ensure no risk of cross-contamination. Cells were washed with PBS once and lysed by adding RIPA buffer supplemented with cOmplete^TM^ mini EDTA-free protease inhibitor cocktail (Roche) directly into the wells followed by incubation on ice for 30 mins. Following centrifugation at 21,300 g for 30 min at 4°C to remove cellular debris, the supernatant was stored at -80°C until further processing. Protein concentration was measured with the Pierce™ BCA Protein Assay Kit (Thermo) following manufacturer’s instructions. To enable the detection and quantification of low levels of tau, total tau was enriched by IP with the polyclonal K9JA anti-tau antibody clone (Dako, cat. no. A0024, Lot No. 20043728) from 400 μg protein lysate for the Exon 1 *MAPT+/+* and *MAPT-/-* #2 lines, 200 μg protein lysate for the Exon 1 *MAPT-/-* #1 line and 450 μg protein lysate for the Exon 4 *MAPT+/+* and *MAPT-/-* lines. For this, protein lysates were incubated in low-protein binding tubes with the antibody (20 μg antibody/mg protein lysate) overnight at 4°C with end-over-end rotation (7 rpm). The antibody-protein complexes were then captured by addition of Protein G magnetic Dynabeads (5 μL beads/μg antibody; Thermo Scientific, cat. no. 13424229) followed by further incubation for 1 h at 4°C with end-over-end rotation. Flow-through containing unbound proteins was removed and beads were washed three times with TBST. Beads were then transferred to fresh low-protein binding tubes and antibody-protein complexes were eluted by incubating in 60 μL 1x Laemmli Sample Buffer (Biorad, cat. no. 1610747, reducing agent was omitted). Eluates were snap-frozen on dry ice and stored at -80°C.

Proteins present in eluates were identified by liquid chromatography-MS/MS analysis. Samples were measured using a Dionex Ultimate 3000 nano-ultra high pressure reversed-phase chromatography system connected to an Orbitrap Fusion Lumos mass spectrometer (Thermo). Peptides were separated by an EASY-Spray PepMap RSLC C18 column (500 mm x 75 μm, 2 μm particle size; Thermo) over a 60 min gradient of 2-35% acetonitrile in 5% DMSO, 0.1% formic acid and at a flow rate of 250 nL/min. The column temperature was maintained at 50°C. The mass spectrometer was operated in positive polarity mode with a capillary temperature of 275°C. Data-independent acquisition mode was utilized for automated switching between MS and MS/MS acquisition, as previously described (23). Approximately 200 ng of peptide material was loaded on column. Samples from the *MAPT-/-* lines were run on the mass spectrometer before those from the *MAPT+/+* lines to eliminate any risk of carry-over. As an additional precaution, blank samples were run on the MS instrument before the eluates from the *MAPT-/-* lines. Data was searched using data-independent acquisition-neural network (version 1.8) against a *Homo sapiens* Uniprot database (retrieved on 16/04/2021), using default settings. All MS data was deposited in Proteome Xchange (PRIDE) and can be accessed using the unique identifier PXD038918.

### Extraction of human brain homogenate

The AD post-mortem brain homogenate extraction protocol was modified from Jin *et al*. (24). To elaborate, 1 g of AD cortical tissue was thawed on ice prior to homogenisation in a Dounce homogeniser for 25 strokes in 4 mL of cold artificial CSF (aCSF; 124 mM NaCl, 2.8 mM KCl, 1.25 mM NaH_2_PO_4_ and 26 mM NaHCO_3_, pH = 7.4) supplemented with a panel of protease inhibitors (5 mM EDTA, 1 mM EGTA, 5 ug/ml leupeptin, 5 µg/ml aprotinin. 2 µg/ml pepstatin, 120 µg/ml Pefabloc and 5 mM NaF). The homogenised mixture then underwent ultracentrifugation at 200,000 g for 110 min at 4°C with a SW-41 Ti rotor (Beckman Coulter) and transferred the supernatant into a Slide-A-Lyzer™ G2 Dialysis Cassettes 2K MWCO (Thermo) in a 100X volume of aCSF without the protease inhibitors swirling for 72 h. The aCSF bath was replaced every 24 h. The resultant brain homogenate was frozen at -80°C.

The AD cortical tissues were obtained from the Oxford Brain Bank. Whole temporal cortices from an AD patient and healthy control (73 years old, female, Braak stage VI, 75 h delay post-mortem, *APOE* ε3/ε3; 70 years old female healthy control, Braak stage I, 81 h delay post-mortem, *APOE* ε3/ε3) were used for this study. In addition, approximately 4 g each of frontal cortices from two AD patients (#1: The same 73- year-old female and #2: 81 years old, male, Braak stage VI, 26 h post-mortem delay, *APOE* ε4/ε4) were used to enrich for AD brain-derived Aβ. The brain homogenates from both frontal cortices were pooled together for experiments.

#### Immunodepletion and enrichment of AD brain-derived Aβ

Immunodepleted (ID) controls were included in a subset of the experiments using AD brain homogenate as Aβ insult. First, Protein A (if the antibody host was rabbit) or Protein G (if the antibody host was mouse) agarose beads (Abcam) were washed three times in aCSF and centrifuged at 400 g for 5 min at 4°C. The beads were then resuspended in 50% slurry with aCSF. At the same time, the brain homogenate aliquots were thawed on ice and centrifuged at 16,000 g at 4°C for 2 min to remove any insoluble component. The agarose beads were then added to the brain homogenate at 3% v/v. Thereafter, 3 mg/ml of S97 anti-Aβ antibody (extracted from a vial of rabbit serum provided by Professor Dominic Walsh; host species: rabbit) at 1:100 dilution (for immunodepletion) or 3 μg/ml of each 4G8 and 6E10 anti-Aβ antibodies (Biolegend) (for brain-derived Aβ enrichment) were added to the brain homogenate-agarose beads mixture. Mock immunodepletion was performed on the AD and healthy control brain homogenates with normal rabbit or mouse IgG antibodies (Abcam).

The mixture was left rotating at 4°C for 12 h before it was centrifuged at 400 g for 5 min at 4°C. The supernatant was transferred for another two rounds of mock or actual immunodepletion whereas the beads were kept in PBS at 4°C for the brain-derived Aβ elution process later. Finally, the supernatant was mixed with Protein A or G agarose beads only (at 2% v/v) for 2 h before the brain homogenate was used for treatments on iPSC-derived cortical neurons.

The antibody-coated agarose beads that pulled down Aβ were washed 3 times with 1 ml of PBS, each time settling the beads by centrifugation at 400 g for 5 min at 4°C. The beads-bound Aβ was then eluted by adding 0.1 M glycine (pH 2.8) that was equivalent to 3 times in volume of the beads. The tube was left rotating at room temperature for 10 min and the eluate was transferred to an equal volume of 1 M Tris (pH 8.0). This elution step was repeated for another two times on the same beads before the eluate was transferred to a centrifuge filter protein concentrator (3 kDa molecular weight cut-off). The concentrators underwent centrifugation at 14,000 g for 2 h at 4°C and the Aβ solution that remained above the filter was diluted in PBS before storing at -80°C.

### Soluble Aβ quantification assay

The levels of Aβ in the brain homogenates were quantified using a V-Plex Plus Aβ Peptide Panel 1 (6E10) Kit from Meso Scale Discovery according to the manufacturer’s instructions. The samples of interest were diluted 1 in 2 in the assay buffer for quantification. The Meso Scale Discovery Workbench 4.0 software was used to analyse Aβ levels.

The AD brain-derived Aβ levels were quantified using a Cisbio Aβ_1-40_ kit. The Aβ levels were detected via homogeneous time-resolved fluorescence from a pair of antibodies. The samples were diluted 1 in 8 for incubation overnight at 4°C. The plate was then read in a PHERAstar^®^ microplate reader (BMG Labtech) to detect fluorescence signals at wavelengths 665 and 620 nm. The data were represented as ΔF which is a relative value to the 665/620 nm signal ratio of the negative control.

### Oligomerisation of synthetic Aβ_1-42_ peptides

Both lyophilised Aβ_1-42_ and treatment control scrambled Aβ_1-42_ peptides (Bachem, H- 1368 and H-7406) were resuspended to 1 mM in Hexafluoro-2-propanol (HFIP). The tubes were vortexed and left sitting at room temperature for 30 min, before they were aliquoted and dried in a Speed-Vac concentrator for 30 min (Thermo) and stored at - 80°C. To oligomerise the Aβ_1-42_ peptides, the Aβ film was first resuspended in dimethyl sulfoxide to 5 mM before it was sonicated in a water bath at room temperature for 10 min. The peptides were subsequently diluted to 100 µM in PBS and the tubes were left stationary at 4°C for 24 h. Just before treating the cells with Aβ oligomers, the solution was centrifuged at 14,000 g for 10 min at 4°C to remove any precipitate/fibrils. The Aβ_1-42_ oligomers were used at a concentration range from 2 to 10 µM.

### Immunocytochemistry

Adherent neurons were fixed in 4% paraformaldehyde for 5 min (for synaptic staining) or 20 min (for other markers), followed by treating with 0.05% saponin in PBS for 20 min for permeabilisation. To block the samples, the plates were treated with 10% normal goat serum (NGS) with 0.01% tween-20 in PBS for 30 min. Primary antibodies were then left incubating with the samples at 4°C overnight with 1% NGS and 0.01% tween-20, before washing 3 times with PBS. Secondary antibodies were then applied in 1% NGS and 0.01% tween-20 at room temperature for 1 h before washing for another 4 times before imaging on the same day. The primary and secondary antibodies used are listed in Supplementary Table 2.

### High-content confocal microscopy imaging

#### Punctate synaptic markers

Neurons seeded in 96-well plates were imaged on a Perkin Elmer Opera Phenix high-content imager. Fifteen images were captured per well with a 43X objective at +1 µm focus level with a binning value of 1. The images were analysed with the Harmony software v4.9 from Perkin Elmer with a customised pipeline. To elaborate, the MAP2-positive neurites were identified with a 0.5-unit overall threshold as the region of interest and resized by expanding outwards by 5 px to cover synaptic signals which lay slightly above the MAP2 signals. Both presynaptic (SYNAPSIN I/II) and postsynaptic (HOMER1) signals were then identified with the Method A of the “Find spots” function with threshold values of 0.17 and 0.14, respectively, with spots larger than 100 px^2^ filtered away. Finally, the synapses were ascertained by finding HOMER1 signals in the vicinity of SYNAPSIN I/II signal regions which had been resized by expanding outwards by 5 px. The absolute number of synapses was then normalised to the total MAP2-positive area to derive synaptic density which was used for all downstream analyses. Synapse loss was achieved by adding the following Aβ insults to the neurons: (1) 10 µM of Aβ_1-42_ oligomers for 24 h, (2) 12.5% (v/v) of AD brain homogenate for 5 days or (3) AD brain-derived Aβ at 200 pg/ml for 5 days.

#### Nuclear staining such as cortical markers and cleaved caspase 3 (CC3)

Fifteen images were captured at -1, 0 and +1 and µm focus levels per well with a 20X objective and binning value of 2. The images were analysed on the same Harmony software by first identifying human nuclei among the co-culture with rat astrocytes and filtering away the nuclei with circularity less than 0.6 units. The percentage of CC3+ cells was calculated by selecting the human nuclei with mean signal intensity greater than a threshold which was determined as the mean intensity across all human nuclei identified plus two standard deviations. To induce cytotoxicity and CC3 overexpression, the neurons were treated with 10 µM of Aβ_1-42_ oligomers for 5 days.

### Live neurite tracing

To accurately trace neurite with live imaging, a low titre of lentivirus expressing both hNGN2 and green fluorescent protein (GFP) was used to transduce NPCs at Day 25 that led to the expression of GFP in approximately 1% of the neuronal population. The hNGN2-eGFP virus was introduced in addition to the regular Ngn2 virus on Day 25 of the cortical neuron differentiation. 0.25 μL (0.025% v/v in the neuronal media) of the hNGN2-GFP virus per well in a 12-well plate per vial of NPCs thawed, and the rest of the differentiation protocol remained the same. The neurons were seeded in co-culture with rat cortical astrocytes on Day 32 at 25,000 cells/cm^2^, and the live imaging session started on Day 35 with the neurons imaged daily for five days. For neurite tracing experiments with exogenous Aβ insults, 10 µM of Aβ_1-42_ oligomers or the scrambled control were added. The whole well was imaged with a 20X objective with 1% overlap between the fields of view to ensure neurite continuity between images.

The images were analysed with the Harmony software with a customised pipeline. Both nuclei and neurites could be identified with the same GFP signal which is stronger at the nuclei. The nuclei were first identified with a “Find nuclei” function with Method B and the identified nuclei with circularity less than 0.6, length greater than 10 µm and width to length ratio greater than 0.3 were filtered away to remove false positives. The neurites were then traced from the nuclei with a “Find neurite” function. Four parameters were selected in this study: (1) Total neurite length per neuron, (2) maximum neurite length per neuron, (3) ramification index per neuron and (4) neurite branch length per branch.

### Cytotoxicity assay (Adenylate kinase)

A ToxiLight^TM^ Non-Destructive Cytotoxicity BioAssay Kit (Lonza) was used to quantify levels of cytotoxicity in neuronal culture supernatant by measuring the release of adenylate kinase (AK). The AK detection reagent was made by mixing the lyophilised reagent with the AK assay buffer supplied in the kit at room temperature. The AK detection reagent was left for 15 min at room temperature shielded from light before 100 µL of the reagent was added to 25 µL of cell supernatant per well for 5 min at room temperature. The plate was then read in a PHERAstar^®^ microplate reader (BMG Labtech) by detecting the levels of fluorescence intensity. Higher fluorescence intensity indicates greater amount of AK released into the neuronal culture supernatant i.e. higher cytotoxicity levels. The AK assay was not performed for the experiments that involved brain homogenate treatments as the soluble factors interfered with the AK detection reagent.

### Multi-electrode array (MEA) assay

The MEA system from Axion Biosystems was used in this study. The iPSC-derived cortical neurons were seeded in the CytoView 48-well MEA plates (16 electrodes per well) on Day 32 as a droplet in the middle of a well to prevent the neurons from coming in contact with the grounding electrodes. The plates were pre-coated in PO/L in a droplet and subsequently 15,000 neurons were seeded per well in a 5 µL droplet. The neurons were allowed to settle for at least 30 min before 5,000 rat astrocytes in 3 µL were added directly into the existing droplet on the plate. Finally, all cells were allowed to settle for at least 1 h before the wells were filled with 200 µL of neuronal medium on the same day. Around Day 90, baseline neuronal firing activities were first measured in the iPSC-derived cortical neurons in a Maestro Pro MEA equipment at 37°C and 5% CO_2_ using the AxIS Navigator^TM^ v2.0.4 by recording for 6 min. The neurons were then treated with the AD brain homogenate at 25% (v/v). Neuronal activities were measured afterwards for 2 min at 6 h, 24 h, 72 h and 120 h post-treatment. The MEA data were analysed in the AxIS Navigator^TM^ and Neuro Metric Tool v2.5.1 software. All recordings were normalised to the baseline of individual well of neurons (e.g., well A1 on Day 1 to 5 of recording was normalised to the recording of well A1 at baseline).

### Live imaging of axonal transport of mitochondria

AXIS^TM^ (Millipore, discontinued in 2018) and XONA microfluidic devices were used to isolate axons and establish directionality for live imaging. Both types of devices had multiple 450-µm long microgrooves between two chambers and the cell bodies were separated by the microgrooves within each chamber. iPSC-derived cortical neurons were seeded at 100,000 cells/cm^2^ according in co-culture with confluent rat astrocytes on one side (soma chamber) of the microfluidic device. The other side of the device (distal chamber) was filled with the same neuronal media (maintained a volume gradient with less volume on the distal side) except that the concentrations for BDNF and NT-3 were ten times higher to encourage axonal outgrowth towards the distal chambers. To label mitochondria, a MitoTracker^TM^ Deep Red FM (Thermo) dye was used at 100 nM in neuronal medium for 3 h at 37°C. The dye was removed by replacing the staining medium with Hank’s balanced salt solution supplemented with 5 mM glucose and 10 mM HEPES. A NIKON Eclipse TE-2000-U fluorescent microscope with a 20X air objective was used for live imaging by taking time lapses that lasted 150 s to capture at least twenty axons per microfluidic device. The automated microscope manoeuvre and imaging parameters were controlled by the Volocity^®^ v6.3.1 (Perkin Elmer) software.

The time course images were then exported as .tiff files to be analysed with Fiji (25) using a TrackMate macro (26) to track motile mitochondria along axons. The Laplacian of Gaussian filter was applied for the spot detector, and the estimated blob diameter was set to be 6 px. The quality threshold was selected to be 5 units and sub-pixel localisation was applied. The number of stationary spots were determined by the median value of all images and the moving spots were determined by subtracting every image in the time course to the median image. A motile mitochondrion was defined by having travelled continuously for at least 10 s and for more than 2 µm in displacement over 150 s of imaging. Motile mitochondria ratio, mean speed, and displacement of mitochondrial axonal transport were selected as readouts of interest.

For the Aβ insult experiments, baseline images were first captured post-staining before treating the neurons with 2 µM Aβ_1-42_ oligomers for 1 h at 37°C and then imaging again post-treatment. For the positive control of axonal transport inhibition, 10 µg/ml of nocodazole was applied instead for 1 h at 37°C.

### Mitochondrial membrane potential assay

The iPSC-derived neurons seeded at 50,000 cells/cm^2^ in monoculture were aged until Day 50. The neurons were then stained with a JC-10 dye (AAT Bioquest) before having their mitochondrial membrane potential measured by detecting signals in two wavelengths emitted by the dye. For the Aβ insult experiments, the neurons were treated with 2 µM of Aβ_1-42_ oligomers for 1 h at 37°C before removing the oligomers and replaced the medium with DMEM basal medium without phenol red (Thermo) with JC-10 added at 1:1,000 dilution. The plate incubated at 37°C for 1 h while shielded from light, before the dye was removed by aspiration and the neuronal culture was replenished with fresh dye-free medium.

To quantify mitochondrial membrane potential, the JC-10-treated plates were read in a PHERAstar^®^ microplate reader (BMG Labtech) by detecting the ratios of fluorescence intensity at wavelengths 520 and 590 nm. Higher 590/520 ratios indicate greater mitochondrial membrane potential with more JC-10 in aggregated form within mitochondria as compared to the monomeric form outside of mitochondria. The plate was read at baseline for three cycles each 13 min apart before adding 10 µM of carbonyl cyanide m-chlorophenyl hydrazone (CCCP) to completely depolarise mitochondrial membrane and the plate was read for another two cycles. The change in mitochondrial membrane potential (Δψ_m_) is defined as the difference in 590/520 fluorescence ratio between the baseline and depolarised mitochondria caused by CCCP.

### Statistical analyses

All data graphing and statistical tests were performed by the GraphPad Prism v9.2.0 software. NS stands for “not significant”. * p < 0.05, ** p < 0.01, *** p < 0.001, **** p < 0.0001 for all statistical analyses. All data were represented as mean ± SEM. For comparisons between two groups, two-tailed Mann-Whitney test was used; for multiple groups, Kruskal-Wallis test was used to compare the groups to the *MAPT+/+* data with Dunn’s multiple comparison correction applied; for multiple dependent variables (for e.g. genotypes and types of treatment), two-way ANOVA was used either compared to the *MAPT+/+* data or the control treatment data within each *MAPT* genotype with Šídák’s (control vs single treatment/time point) or Dunnett’s (control vs multiple treatments/time points) multiple comparison correction applied. Outlier removal specifically for the axonal transport of mitochondria data was conducted with a ROUT method with 5% false discovery rate applied.

## Results

### Generation and validation of the isogenic *MAPT-/-* iPSC panels

To generate *MAPT-/-* iPSC lines, two parental iPSC lines from healthy individuals were used with two different targeting strategies as described in *Methods* – either a single gRNA targeting *MAPT* Exon 1 in one of the parental iPSC lines, or a pair of gRNAs targeting Exon 4 in the other, to diversify targeting options for achieving successful homozygous CRISPR-Cas9-mediated *MAPT* knockout. The intended edit was either a double-stranded break towards the end of Exon 1 3-bp upstream from the 3’-end of the gRNA binding region, or a 25-bp deletion towards the beginning of Exon 4 flanked by the pair of gRNAs used (Fig. 1A and Supplementary Table 1). After successive rounds of sequencing validations, we identified two *MAPT-/-* iPSC clones from ninety-six clones (2.1% successful targeting efficiency) of the Exon 1-targeted line and one *MAPT-/-* iPSC clone from seventy-two clones (1.4% successful targeting efficiency) of the Exon 4-targeted line. We also identified *MAPT+/-* (heterozygous knockout) and *MAPT+/+* (clones that failed to have their *MAPT* genetically edited and remained as wild-type) from each parental line to be included in downstream experiments. These isogenic iPSC lines with a range of *MAPT* genotypes are herein referred to as Exon 1 or Exon 4 isogenic panels.

**Figure 1:**
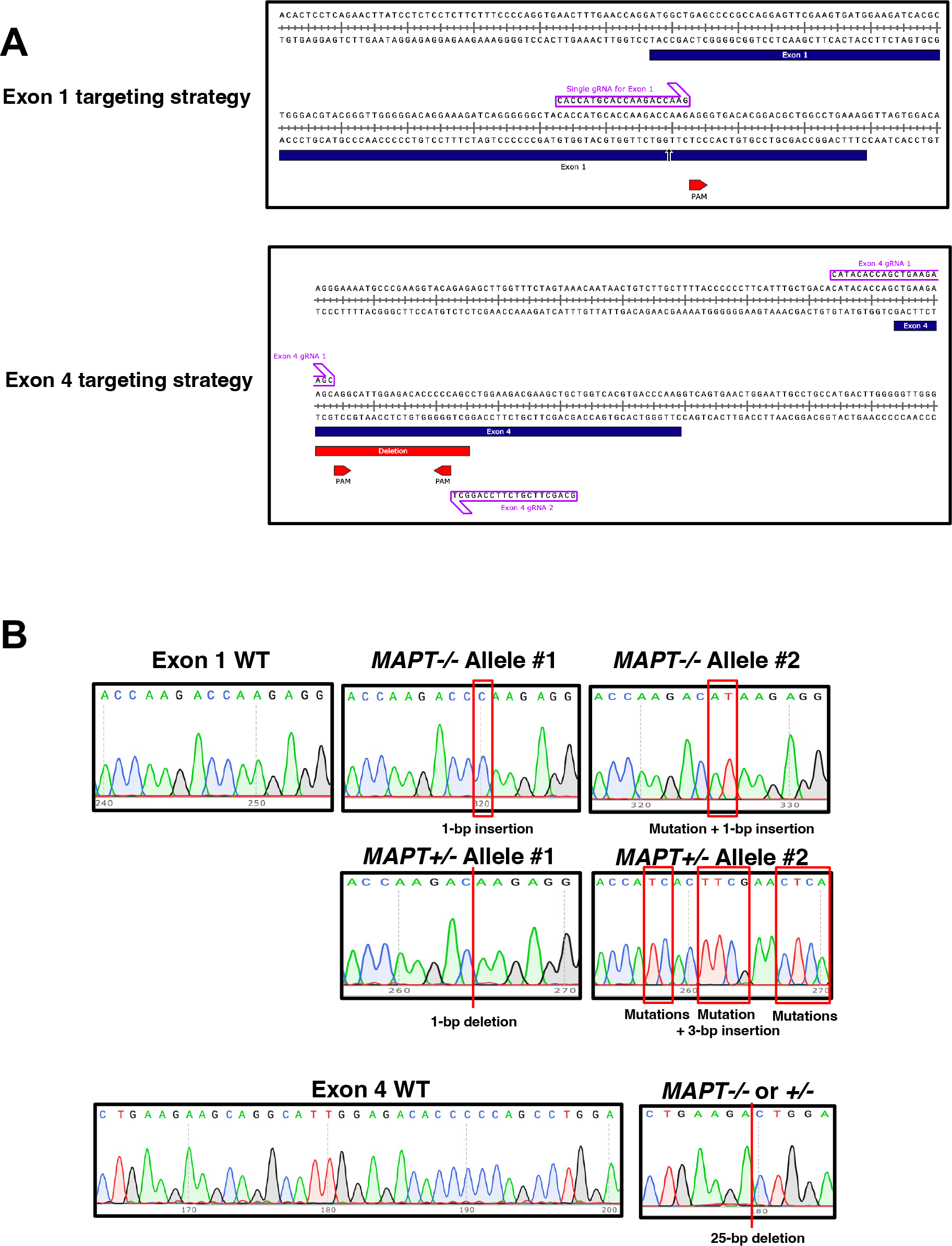
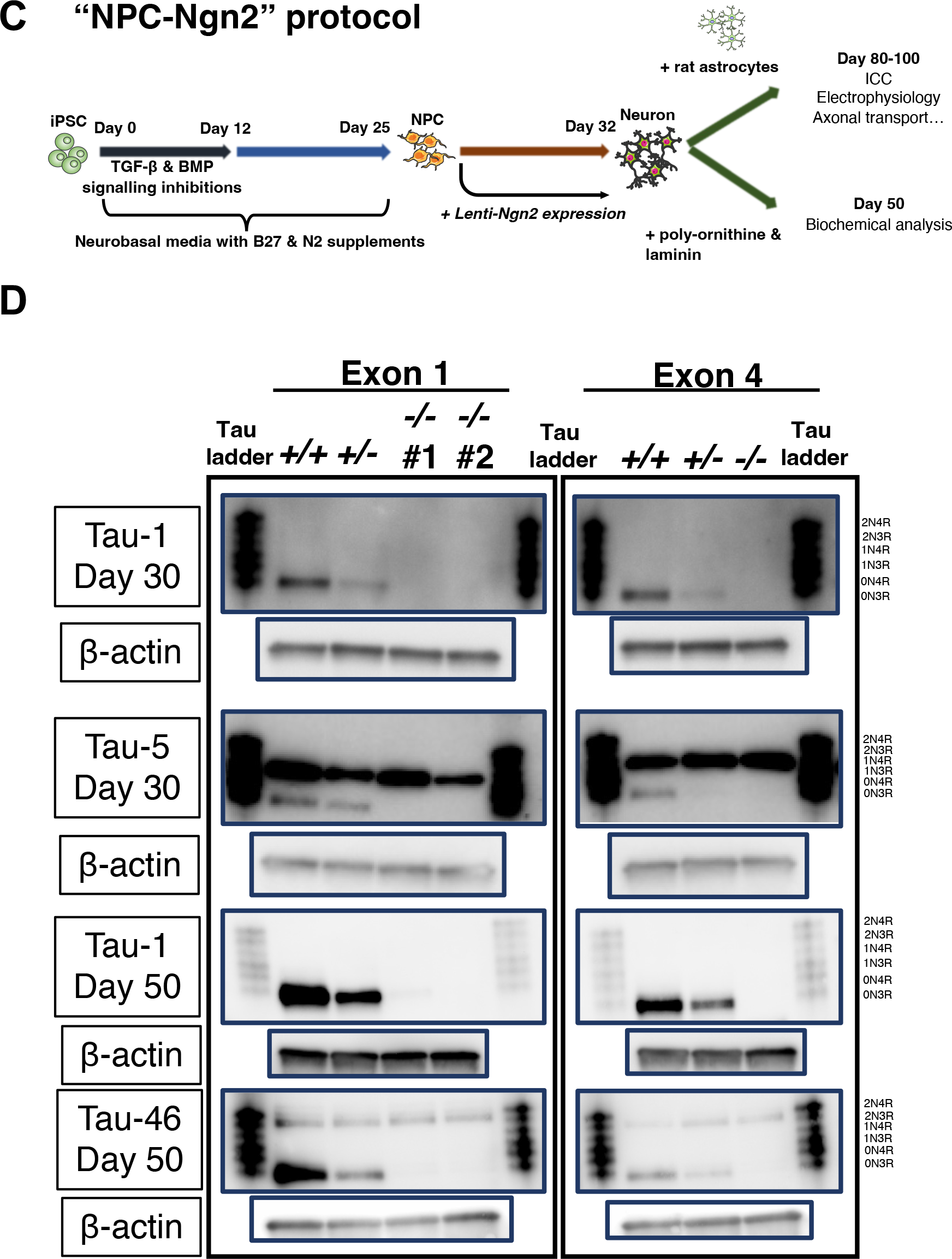
Generation and validation of *MAPT-/-* iPSCs. **(A)** Illustrations of gene editing strategies showing the positions of gRNA(s) in Exon 1 and 4 DNA loci. The intended double-stranded break in Exon 1 is indicated by an arrow, whereas the intended 25-bp deletion is indicated between the gRNA pair used to target Exon 4. **(B)** Sequencing results of edited *MAPT* loci, including all combinations of alleles in the Exon 1 isogenic panel. For the Exon 4 isogenic panel, both *MAPT+/-* and *MAPT-/-* lines harbour the 25-bp deletion at the same site. **(C)** Schematic of the cortical neuron differentiation protocol used throughout this study– iPSC lines were first differentiated concurrently to NPCs, before they were subject to lentiviral transduction for *Ngn2* expression (“NPC-Ngn2 protocol”) for maturation either in co-culture with primary rat cortical astrocytes or in monoculture on coated surface. **(D)** Western blots probing for tau using three different antibodies (Tau-1 – mid-region, Tau-5 – mid-region and Tau-46 – C-terminus) either on Day 30 (NPC) or Day 50 (neuron) of neuronal differentiation for both *MAPT-/-* isogenic panels. 6 ng of recombinant tau ladder was used and 5 μg of lysate was added per lane (except for the Tau-46 blot where 10 μg of lysate was added). Anti-β-actin blots were used as the expression control housekeeping protein for the lysates.

The Exon 1-targeting strategy produced a range of genetic alterations due to a single DNA double-stranded break. In one of the DNA sequencing-confirmed Exon 1 *MAPT-/-* lines (*MAPT-/-* #1), one allele carries a 1-bp insertion whereas the other allele has a single-nucleotide mutation in addition to the insertion. Both genetic alterations resulted in a single-nucleotide frame shift which gave rise to a stop codon within Exon 1 (Fig. 1B; *MAPT-/-* Alleles #1 and #2). The other Exon 1 *MAPT-/-* line (*MAPT-/-* #2) has a single-nucleotide insertion as in *MAPT-/-* Allele #1 in both alleles. In the Exon 1 *MAPT+/-* line, one of the alleles sustained a 1-bp deletion (*MAPT+/-* Allele #1), similarly giving rise to a frame shift and a stop codon within Exon 1. The other allele experienced a 3-bp in-frame insertion, coupled with several mutations around the insertion site to causing changes in five amino acid residues without producing a stop codon (*MAPT+/-* Allele #2). The Exon 4 *MAPT-/-* and *MAPT+/-* lines both carry the same expected 25-bp deletion that led to a frame shift plus a stop codon within Exon 4 (Fig. 1B).

We subsequently differentiated the Exon 1 and 4 isogenic panels into Day 30 NPCs and Day 50 cortical neurons as described in *Methods* and illustrated in Fig. 1C, before collecting cell lysates for western blots for probing for tau proteins. Antibodies targeting either the mid-region (Tau-1 from amino acid 192 to 204 and Tau-5 from amino acid 218 to 225) or the C-terminus (Tau-46 from amino acid 404-441) of tau did not detect any 0N3R tau isoform which would be the only isoform readily expressed by the iPSC-derived NPCs and cortical neurons on Day 30 and 50, respectively (Fig. 1D). A band corresponding to the 2N3R tau isoform was also noticeable in Tau-5 and Tau-46 blots across all lines, but the band was considered unspecific. Since tau isoform expression is developmentally regulated (15) i.e., expressing progressively from the shortest isoform in foetus to eventually including the full-length 2N4R tau isoform in adults, it is unlikely that the cells were expressing a more mature tau isoform without expressing the 0N3R isoform first. Moreover, the iPSC-derived neurons up to 50 days into differentiation are relatively immature neuronal cells thus unlikely to be expressing mature tau isoforms with N-terminus inserts at high levels (27).

We found, however, that by extending the Western blot exposure time as well as saturating the chemiluminescence signals detected for the *MAPT+/+* lines, a specific band with a molecular weight lower than that of the 0N3R (smallest) isoform became detectable in both the *MAPT+/-* and *MAPT-/-* lines from the Exon 4 panel especially in the relatively more mature Day 50 neurons (Supplementary Fig. 2). As this tau-immunoreactive band was not present in the *MAPT+/+* line, we reasoned that it may represent a non-canonical protein product of the *MAPT* gene that was generated in response to the out-of-frame deletion in Exon 4. Moreover, we noticed a faint band at the molecular weight corresponding to the 0N3R isoform position in the Exon 1 *MAPT-/-* #1 specifically in the Day 50 neurons (Tau-1 blot) that was undetectable in the original blot with the settings required to detect physiological levels of tau expression in the *MAPT+/+* neurons. We then further tested tau expression in Day 50 iPSC-derived cortical neurons from both isogenic panels by subjecting the cell lysates for immunoprecipitation-mass spectrometry (IP-MS) analysis. The IP-MS results indicated that tau peptides were indeed detectable in all *MAPT-/-* lines (Supplementary Fig. 3A) but the MS signal intensity (an estimate for quantity) was very low and significantly less than that in the *MAPT+/+* neurons at 1%, 0.06% and 11% for the Exon 1 *MAPT-/-* #1, Exon 1 *MAPT-/-* #2 and Exon 4 *MAPT-/-* neurons, respectively (Supplementary Fig. 3B). This IP-MS experiment was unable to identify the specific non-canonical tau peptide observed in the Exon 4 *MAPT+/-* and *MAPT-/-* lines, thus it is unclear if those tau peptides are functional. We therefore concluded that these *MAPT-/-* lines retain residual tau expression, but the expression levels are extremely low for the Exon 1 panel with an estimated ≤1% tau expression, an almost total depletion, whereas the Exon 4 *MAPT-/-* neurons expressed a non-canonical tau peptide level at approximately 11% relative to the *MAPT+/+* neurons.

### Tau depletion protects neurons from AD brain-derived Aβ-driven hyperactivity

The Exon 1 and Exon 4 isogenic panels were then used for downstream experiments detailed in this study (Supplementary Table 3). We began by examining relatively more sensitive phenotypes such as neuronal activity and synapse loss, to axonal transport of mitochondria that involves a shorter time scale before examining relatively more severe cellular phenotypes such as neurite outgrowth impairment and neurodegeneration. To address if tau depletion affects neuronal activity, we differentiated the *MAPT+/+* and *MAPT-/-* lines from both Exon 1 (*MAPT-/-* #1) and Exon 4 isogenic panels into cortical neurons (Supplementary Fig. 4) on MEA multi-well plates which have sixteen electrodes embedded at the bottom of the plates per well for extracellular field potential detection (Fig. 2A). The *MAPT-/-* neurons from both Exon 1 and 4 panels exhibited marked impairments in neuronal activity across all parameters quantified i.e., showing reduced firing strength, frequency, synchronicity within the neuronal network and network firing (Fig. 2B).

**Figure 2:**
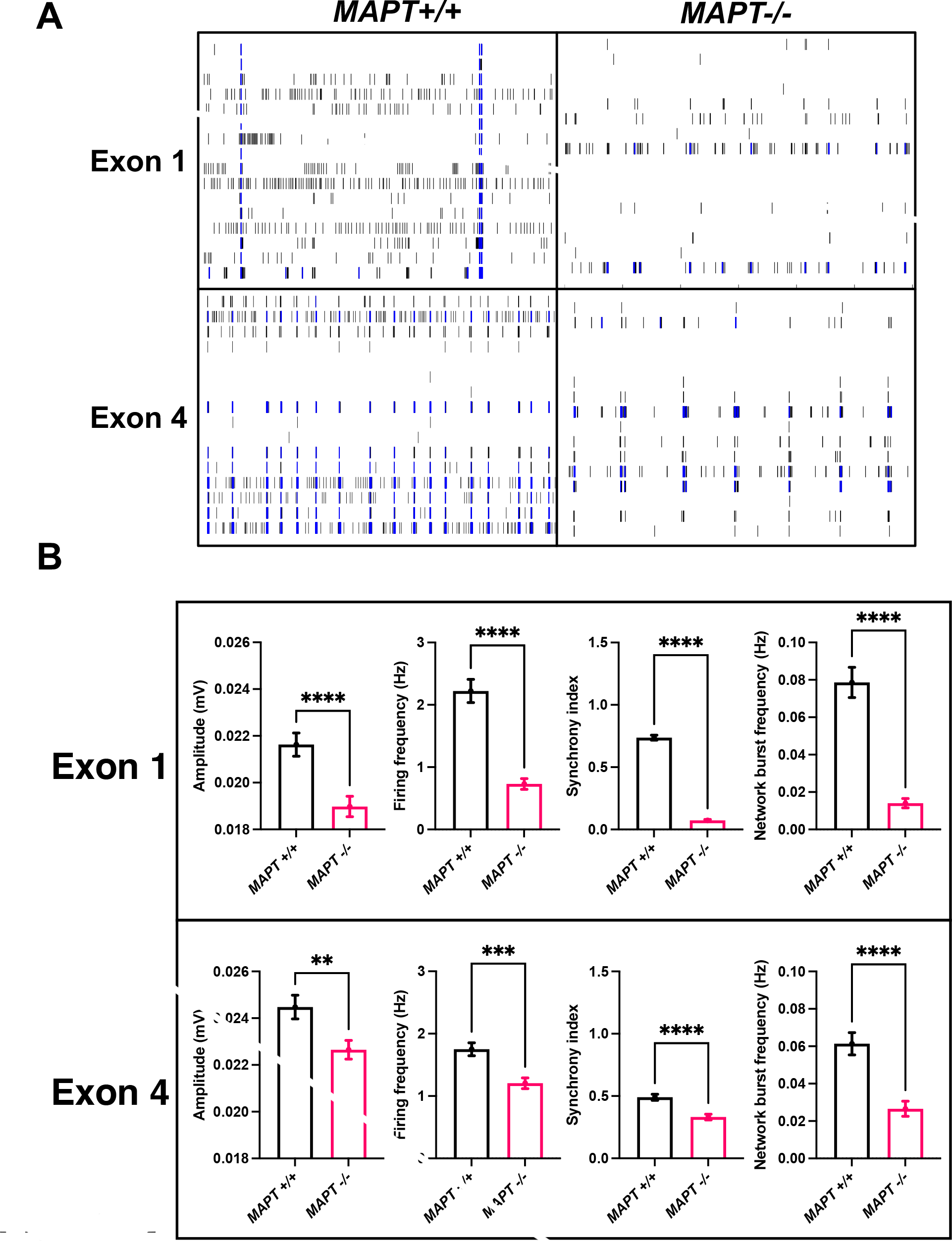
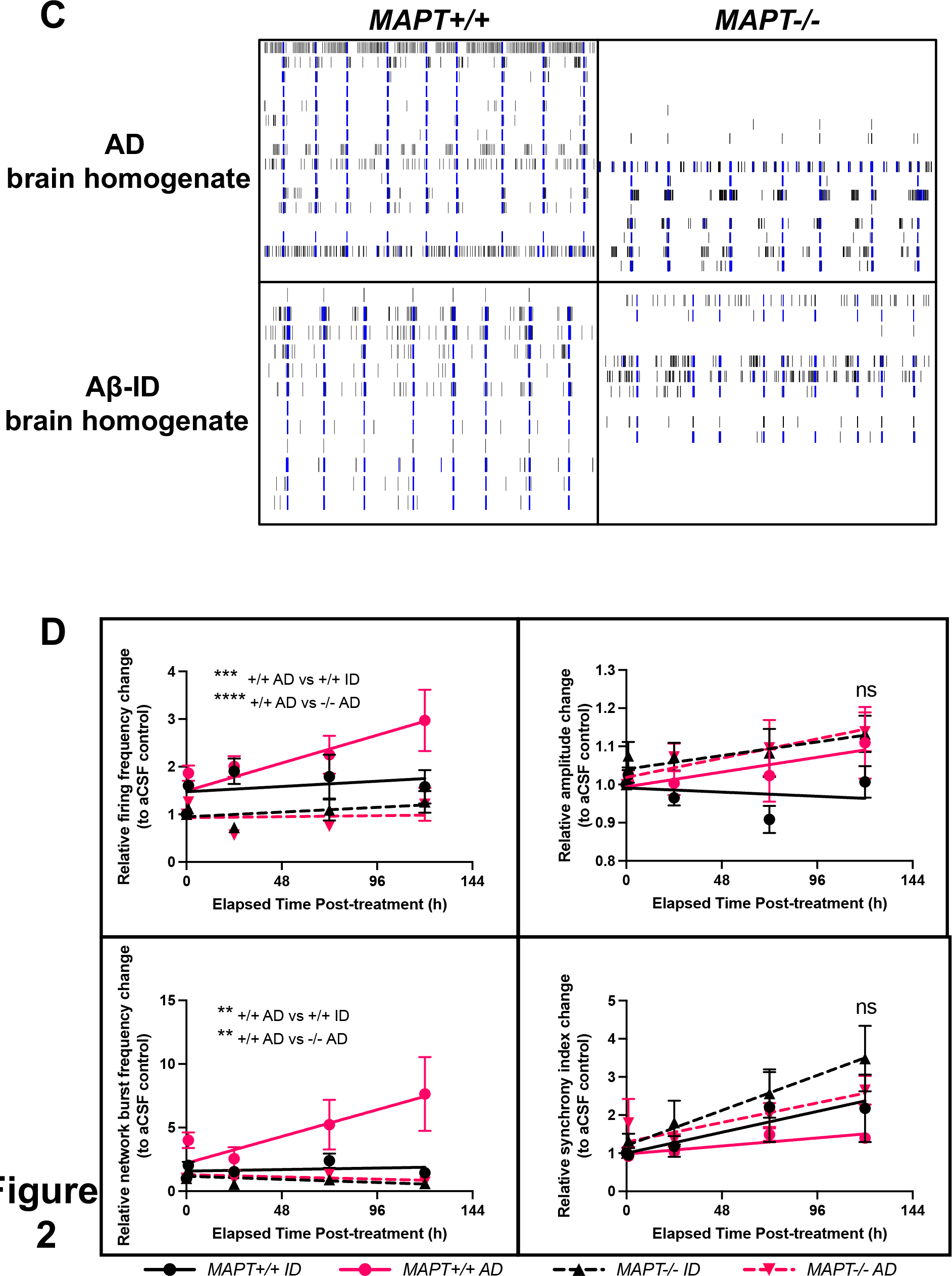
*MAPT-/-* iPSC-derived cortical neurons demonstrate reductions in neuronal activity and protection from Aβ-driven hyperactivity. **(A)** Representative raster plots showing individual neuronal activity spikes for each of the sixteen electrodes (row) in each MEA well over 2 min for both Day 90 *MAPT+/+* and *MAPT-/-* neurons (*MAPT-/-* #1 from the Exon 1 panel) from each isogenic panel at baseline. **(B)** Quantification of baseline neuronal activity parameters measured by MEA assays on Day 90-100 iPSC-derived cortical neurons from both *MAPT-/-* isogenic panels. Mean ± SEM. *n* = 53-55 (Exon 1 *MAPT+/+*) or 51-53 (Exon 1 *MAPT-/-*) wells across three independent neuronal differentiation repeats; 101-105 (Exon 4 *MAPT+/+*) or 103-114 (Exon 4 *MAPT-/-*) wells across six independent neuronal differentiation repeats. Some wells did not achieve the threshold needed to register network activities. Two-tailed Mann-Whitney test was used for statistical analysis. **(C)** Representative raster plots showing individual neuronal activity spikes for each of the sixteen electrodes (row) in each MEA well over 2 min for both Day 90 Exon 4 *MAPT+/+* and *MAPT-/-* neurons treated with either AD brain homogenate or Aβ- immunodepleted (ID) AD brain homogenate at 25% v/v in the neuronal media for 5 days. **(D)** Quantification of neuronal activity parameters measured by MEA assays over 5 days on Day 90-93 Exon 4 *MAPT+/+* and *MAPT-/-* iPSC-derived cortical neurons treated with either AD brain homogenate or Aβ-ID homogenate at 25% v/v in the neuronal media. All datapoints were normalised to the baseline recording pre- treatment, and for each time point relative to the wells subject to aCSF (vehicle) control treatment. Mean ± SEM. *n* = 7-14 (*MAPT+/+* ID), 11-14 (*MAPT+/+* AD) or 5-16 (*MAPT-* */-* ID and AD) wells across three independent neuronal differentiation repeats. Two- way ANOVA with Dunnett’s multiple comparison correction was used for statistical analysis compared against the *MAPT+/+* AD wells 5 days post-treatment.

We then asked whether tau lowering can protect human iPSC-derived cortical neurons from exogenous toxic insults such as Aβ by treating neurons with AD brain homogenate, or with AD brain homogenate immunodepleted for Aβ (Aβ-ID) and assaying neuronal activity by MEA (Fig. 2C). The *MAPT+/+* line demonstrated AD brain homogenate-driven hyperactivity over time both in terms of single-electrode neuronal firing and network firing frequencies, whereas neuronal firing amplitude and synchrony remained unaffected (Fig. 2D). This hyperactivity phenotype was not present after treatment by Aβ-ID, or after tau depletion, suggesting that the hyperactivity phenotype was specifically Aβ-driven and tau-dependent.

On the synapse level, tau depletion did not affect synaptic density in the iPSC-derived cortical neurons (Supplementary Fig. 5A and 5B). Bulk AD brain homogenate treatment led to a 20-30% synapse loss in the iPSC-derived cortical neurons regardless of their *MAPT* genotypes or the presence of Aβ (Supplementary Fig. 5C; Exon 4). We then performed extraction and concentration of Aβ from the AD brain homogenate as detailed in *Methods* and showed that the treatment of AD brain- derived Aβ resulted in approximately 10% synapse loss in the *MAPT+/+* neurons but not in the *MAPT+/-* and *MAPT-/-* neurons (Supplementary Fig. 5D; Exon 4). This indicates that AD brain-derived Aβ-driven synapse loss is tau-dependent, but that there are other soluble factors present in the AD brain homogenate that can result in synapse loss. Both bulk AD brain homogenate and brain-derived Aβ treatments were insufficient to cause synapse loss in the Exon 1 isogenic panel (Supplementary Fig. 5C and 5D; Exon 1).

### Tau depletion mitigates Aβ-driven deficit in retrograde axonal transport of mitochondria

Since tau is mainly localised in the axons as a microtubule-binding protein (28), we next asked whether tau depletion interferes with axonal transport of mitochondria. The iPSC-derived cortical neurons were plated in one side of microfluidic chambers for live imaging of mitochondrial movement along axons with clear directionality as detailed in *Methods* (Fig. 3A). We did not observe any changes in the ratio (to stationary mitochondria), speed and displacement of motile mitochondria in the *MAPT-/-* neurons at baseline (Supplementary Fig. 6A).

**Figure 3:**
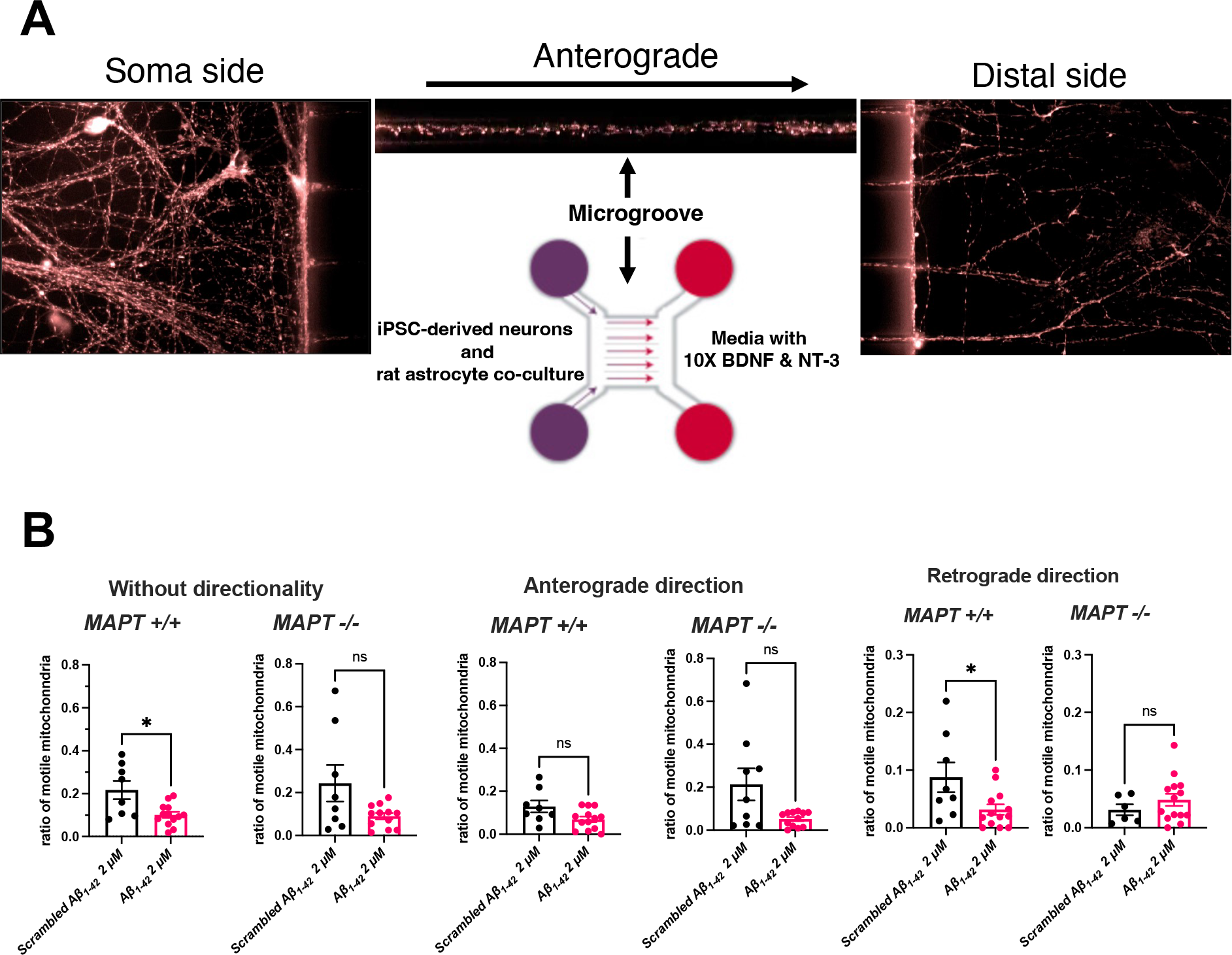
Aβ-driven retrograde impairment of axonal transport of mitochondria is absent in *MAPT-/-* iPSC-derived cortical neurons. **(A)** Schematic of the experiments designed to measure axonal transport of mitochondria using microfluidic chambers. **(B)** Quantification of ratio of motile mitochondria (motile to stationary) in Day 70-95 iPSC-derived cortical neurons from the Exon 4 isogenic panel with or without directionality over 150 s of live imaging. The neurons were treated with either 2 μM scrambled Aβ_1-42_ or Aβ_1-42_ oligomers for 1 h before imaging. Mean ± SEM. *n* = 8 (*MAPT+/+* scrambled Aβ_1-42_), 13 (*MAPT+/+* Aβ_1-42_), 6-9 (*MAPT-/-* scrambled Aβ_1-42_) and 12-14 (*MAPT-/-* Aβ_1-42_) microfluidic chambers measured across five independent neuronal differentiation repeats. Two-tailed Mann-Whitney test was used for statistical analysis.

We subsequently investigated whether tau depletion mitigates neuronal response to Aβ in terms of impairments to axonal transport of mitochondria. Recombinant Aβ_1-42_ oligomers were used as a source of a more acute and toxic Aβ insult to the neurons, as previously reported in mouse primary hippocampal neurons *in vitro* (8). After one hour of exposure to Aβ_1-42_ oligomers, there were fewer motile mitochondria in the *MAPT+/+* neurons specifically in the retrograde direction while the Aβ-driven reduction in the number of mitochondria moving towards the soma was mitigated in the *MAPT-/-* neurons (Fig. 3B). This suggests that the neuronal response to exogenous Aβ insult in axonal transport, at least for mitochondria as a cargo, is tau dependent. The treatment of exogenous Aβ_1-42_ oligomers did not result in any changes in either speed or displacement of motile mitochondria along axons in the iPSC-derived cortical neurons regardless of their *MAPT* genotypes (Supplementary Fig. 6B). We also found that there were no differences in mitochondrial membrane potential within the Exon 1 and 4 isogenic panels at basal level, nor in response to exogenous Aβ_1-42_ oligomers, indicating that the tau-dependent phenotype in axonal transport of mitochondria observed was unrelated to mitochondrial function (Supplementary Fig. 6C and 6D).

### Tau depletion does not consistently result in neurite outgrowth impairments

We went on to investigate whether tau lowering can result in changes in neuronal morphology. To address this question, we transduced a subset of iPSC-derived cortical neurons with vectors expressing GFP alongside *NGN2* and tracked their individual neurite outgrowth with live imaging over time (Supplementary Fig. 7A). Both *MAPT-/-* (#1 line) neurons from the Exon 1 isogenic panel and *MAPT-/-* from the Exon 4 isogenic panel exhibited neurite outgrowth impairment with shorter neurite and axonal lengths, as well as lower ramification index (levels of branching per root from soma) as compared to the respective *MAPT+/+* lines without alterations in neurite branch length (Supplementary Fig. 7B). However, the *MAPT-/-* (#2 line) neurons from the Exon 1 isogenic panel did not suffer from any neurite outgrowth impairment and surprisingly demonstrated a reduction in neurite branch length as compared to the *MAPT+/+* neurons suggesting that tau depletion alone is inadequate to consistently result in neurite outgrowth impairments.

### Tau depletion protects neurons from Aβ-driven neurodegeneration

Finally, we asked whether tau lowering can mitigate Aβ-driven neurodegeneration in human iPSC-derived cortical neurons. We again used recombinant Aβ_1-42_ oligomers as a more acute and toxic source of Aβ to induce neurodegeneration in the iPSC-derived cortical neurons and measured the percentage of cleaved caspase 3-positive (CC3+) neurons as a readout for cell death (Fig. 4A). In both Exon 1 and 4 isogenic panels, there was a substantial increase in Aβ_1-42_ oligomer-driven neurodegeneration in the *MAPT+/+* lines (Fig. 4B). Tau depletion was effective in mitigating Aβ_1-42_ oligomer-driven neurodegeneration in both *MAPT+/-* and *MAPT-/-* neurons, a finding that was consistent in both isogenic panels, indicating that this phenotype is tau-dependent and, crucially, that partial tau reduction was also sufficient to mitigate Aβ_1- 42_ oligomer-driven neurodegeneration.

**Figure 4:**
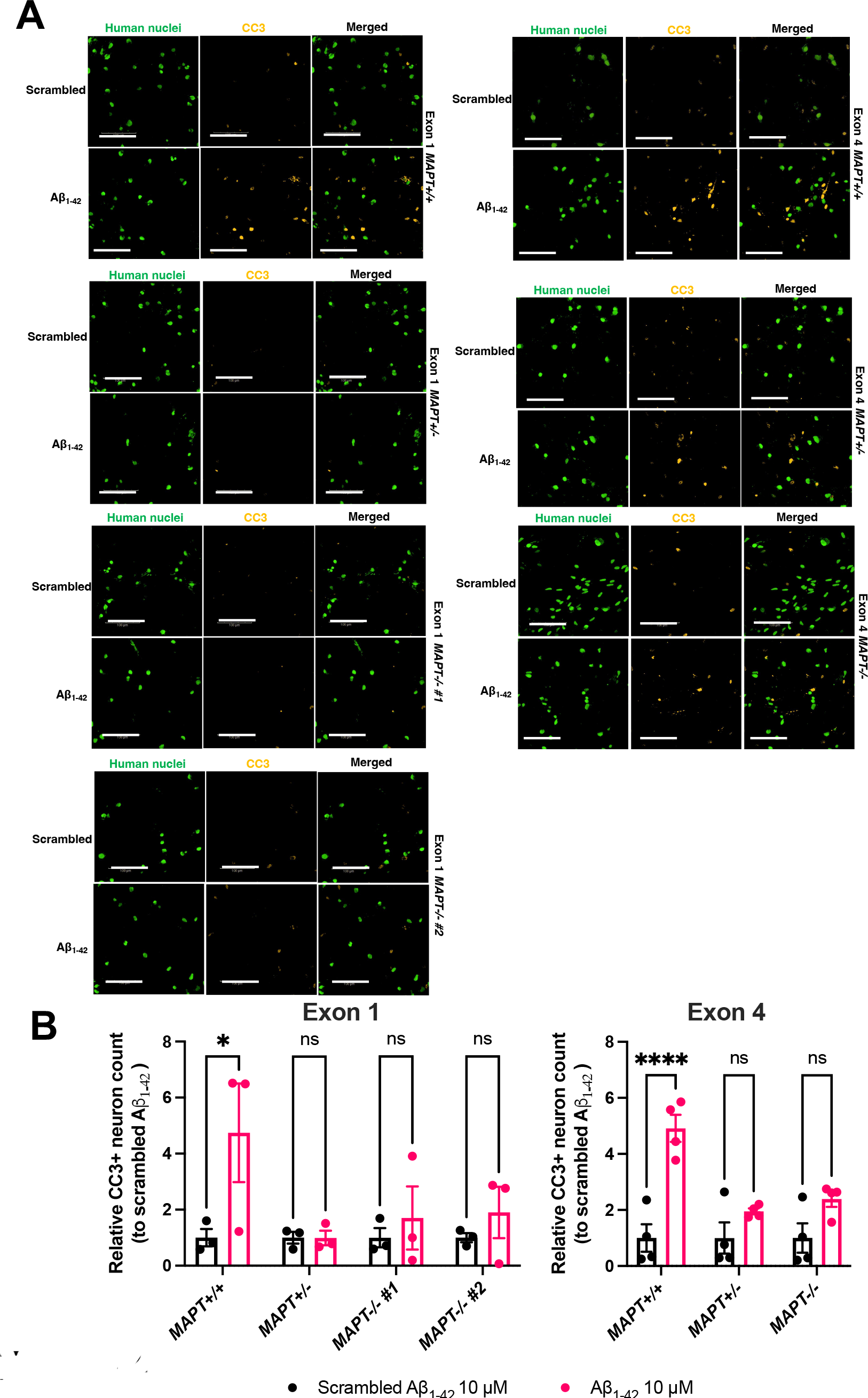
Aβ-driven neurodegeneration is absent in *MAPT-/-* iPSC-derived cortical neurons. **(A)** Representative immunofluorescence images of Day 79-83 (Exon 1 isogenic panel) and Day 79-86 (Exon 4 isogenic panel) iPSC-derived cortical neurons treated with either 10 μM scrambled Aβ_1-42_ or Aβ_1-42_ oligomers for 5 days. The neurons were probed with antibodies against human nuclei (green) and cleaved caspase 3 (CC3; yellow) which was used as the marker for cell death. Scale bar = 100 μm. **(B)** Quantification of relative CC3+ neuron count post-treatment with either 10 μM scrambled Aβ_1-42_ or Aβ_1-42_ oligomers for 5 days. Mean ± SEM. *n* = three (Exon 1) or four (Exon 4) independent neuronal differentiation repeats. Two-way ANOVA with Šídák’s multiple comparison correction was used for statistical analysis compared against the scrambled Aβ_1-42_ treatment.

## Discussion

In this study, we demonstrated that tau depletion in human iPSC-derived cortical neurons caused considerable reductions in neuronal activity without affecting synaptic density. We also observed neurite outgrowth impairments in two of the *MAPT-/-* lines studied. Other phenotypes, including axonal transport of mitochondria, mitochondrial function, and cortical neuron differentiation efficiency remained unaffected by tau depletion. These observations are consistent with *in vivo* studies conducted in *Mapt-/-* mice where the absence of tau did not lead to overt behavioural or cognitive deficiencies and mouse *Mapt-/-* primary neurons exhibited reduced baseline activity and neurite outgrowth impairment *in vitro* (13, 29). Challenging iPSC-derived cortical neurons with exogenous Aβ resulted in hyperactivity, retrograde axonal transport deficit of mitochondria and neurodegeneration but, crucially, we find these adverse effects were mitigated by tau depletion.

In the presence of exogenous Aβ human iPSC-derived cortical neurons experienced hyperactivity which is tau-dependent. This is in line with clinical observations where pre-symptomatic AD patients present with hippocampal hyperactivity, as well as preclinical findings in AD mouse models where neurons in the vicinity of Aβ plaques are hyperactive, a behavioural phenotype that has been found to require tau (30–32). It has also been reported that *Mapt-/-* neurons were less prone to hypersynchrony caused by over-excitation and that *Mapt-/-* mice were protected from Aβ-driven seizure episodes (10, 29). Overall, the diminished baseline neuronal activity seen in both mouse and human tau-depleted neurons may be considered a protective mechanism, rather than a functional deficit.

One of the strengths of our study is the generation and analysis of multiple *MAPT-/-* lines generated from different individuals and targeted using different strategies. Generating multiple lines allowed us to observe line-to-line variation in the response of two iPSC lines derived from different individuals to exogenous Aβ insults. Specifically, the Exon 1 isogenic panel appeared more resilient to the Aβ insults (Fig. 4B, Supplementary Fig. 5C and 5D) highlighting the importance of isogenic controls to eliminate from downstream experiments any effects arising from genetic variations between different individuals. Even within respective isogenic panels we still observed differences between the lines for certain phenotypes, such as neurite outgrowth impairment shown in one of the two Exon 1 *MAPT-/-* lines. This inconsistency in neurite outgrowth has also been observed in different *Mapt-/-* mouse strains and it has been suggested that compensatory expression from other microtubule-associated proteins could potentially account for the difference, although it is unclear how this is regulated in different *Mapt-/-* mouse strains (13, 33).

A challenge in any study modelling AD *in vitro* is the choice of exogenous Aβ insult applicable across a range of phenotypic assays. We found AD brain homogenate to be a physiological source of pathologically relevant Aβ for electrophysiological and synaptic assays with a concentration of Aβ in the pM range. AD brain homogenate had been routinely used in other electrophysiology studies and we found that the hyperactivity readout in human iPSC-derived cortical neurons was particularly sensitive to and specifically caused by the presence of Aβ. However, we did not observe such specificity in the synapse loss experiment, suggesting that other soluble factors present in the AD brain homogenate could similarly lead to synapse loss (Supplementary Fig. 5C). Treating neurons with purified AD brain-derived Aβ demonstrated that the Exon 4 *MAPT-/-* neurons were protected from Aβ-driven synapse loss, although 200 pg/ml of AD brain-derived Aβ was insufficient to cause synapse loss in the Exon 1 isogenic panel (Supplementary Fig. 5D). However, the use of AD brain-derived Aβ is limited by the capture-elution efficiency of the current method in which 1 g of AD brain tissue was needed to extract 1 ng of AD brain-derived Aβ.

Recombinant Aβ_1-42_ oligomers provide a higher concentration of exogenous Aβ resulting in more robust phenotypes. Retrograde axonal transport of mitochondria was impaired by 2 μM of Aβ_1-42_ oligomers in iPSC-derived cortical neurons, consistent with a previous study in mouse primary hippocampal neurons with a similar experimental setup (8). However, in that previous study axonal transport was impaired in both directions and it was later reported that naturally occurring Aβ peptides in an AD mouse model led to axonal transport deficits only in the anterograde direction (34). The consensus between the present human cell model study and previous mouse studies is that Aβ drives a reduction in the number of motile mitochondria transported along axons, although specific effects of Aβ may be dependent on the experimental setup, source of Aβ and/or species of cellular model.

To cause a robust neurodegeneration cell death phenotype, we treated iPSC-derived cortical neurons with 10 μM of Aβ_1-42_ oligomers for five days. This supraphysiological insult consistently resulted in at least two-fold increase in the percentage of CC3- positive neurons, even in the Exon 1 isogenic panel which was more resilient to Aβ challenges (Fig. 4). The *MAPT+/-* and *MAPT-/-* neurons from both isogenic panels consistently demonstrated protection from Aβ-driven neurodegeneration in line with a previous study in *Mapt-/-* primary hippocampal neurons in which 20 μM of Aβ_1-40_ fibrils was used to treat neurons for four days (9). This critical phenotype highlights consistent protective effects of tau lowering in Aβ-driven toxicity in AD pathogenesis in human and mouse cells.

Another strength of our study was to investigate tau expression levels in *MAPT-/-* lines with very sensitive IP-MS methodology in addition to western blots (Supplementary Fig. 2 and 3). We found that the Exon 1 *MAPT-/-* lines carrying single nucleotide insertions expressed extremely low levels (≤1%) of tau peptides as compared to the *MAPT+/+* neurons, an almost total depletion. The Exon 4 *MAPT-/-* line (carrying a 25- bp deletion) produced a non-canonical tau-immunoreactive band of a molecular weight lower than that of the smallest 0N3R tau isoform which likely arose through skipping of exon 4, an event that was previously reported and would neither introduce an early stop codon nor alter the sequence of the remainder of the tau protein (35), at ∼11% of overall expression levels as compared to the isogenic *MAPT+/+* neurons. The IP-MS analysis returned a total of fifty-one peptides from a subset of *MAPT* exons and we were unable to determine the identity of the non-canonical tau peptide seen in the Exon 4 *MAPT-/-* line. Future work may be able to characterise this variant through long-read sequencing of mRNA product(s). Nevertheless, marked tau depletion was achieved in the Exon 4 *MAPT-/-* lines and all phenotypes reported in the Exon 1 *MAPT-/-* lines were consistently reproduced in the Exon 4 *MAPT-/-* line, suggesting the functional effects observed were specifically caused by chronic tau depletion.

## Conclusions

In this study, we established stable human iPSC isogenic panels with chronic tau depletion from two healthy individuals. We demonstrated that a wide range of Aβ- driven phenotypes in iPSC-derived cortical neurons were tau-dependent, including Aβ-driven hyperactivity, axonal transport deficits, and neurodegeneration, consistent with studies conducted in *Mapt-/-* mouse models. Our findings highlight the potential benefits of ongoing attempts at chronic tau-lowering strategies in AD in the clinic. iPSC-derived *MAPT-/-* human cortical neurons can be applied to investigate the involvement of tau in Aβ-driven toxicity in cortical neurons and in other tauopathy-relevant pathways.

## List of abbreviations

aCSF – artificial cerebrospinal fluid; AD – Alzheimer’s disease; Aβ - amyloid-β; BSA – bovine serum albumin; CRISPR – clustered regularly interspaced short palindromic repeats; CC3 – cleaved caspase 3; carbonyl cyanide m-chlorophenyl hydrazone; iPSC induced pluripotent stem cells; ID – immunodepletion; IP – immunoprecipitation; MEA – multiple electrode array; MS – mass spectrometry; PBS – phosphate-buffered saline; PO/L – poly-L-ornithine and laminin; RIPA – radioimmunoprecipitation assay; SDS – sodium dodecyl sulfate; TBST – tris-buffered saline with tween; NIM – neural induction medium; NMM – neural maintenance medium; NPC – neural progenitor cell; NGS – normal goat serum.

## Ethics approval and consent

The ethics information for the SBAd-03-01 parental iPSC line can be found from the Lonza product page (CC-2511) where the source dermal fibroblast was obtained, whereas that for the SFC856-03-04 parental iPSC line was approved by the National Health Service, Health Research Authority, National Research Ethics Service Committee South Central, Berkshire, United Kingdom, Research Ethics Committee 10/H0505/71.

## Consent for publication

Not applicable.

## Data and material availability

The datasets used and/or analysed during the current study are available from the corresponding author on reasonable request. The iPSC lines used in this study can be requested from the James and Lilian Martin Centre for Stem Cell Research.

## Authors’ contributions

B.N. conducted all functional experiments, analysis and material acquisition described in this study. J.V. and S.A.C. conceived the CRISPR-Cas9-mediated gene editing design and generated the *MAPT-/-* iPSC lines. D.B.-K. conceptualised and supervised the electrophysiological experiments. M.I.S. conducted the IP experiment, supervised by J.A.T.. D.P.O. conducted the MS experiment and its analysis. N.B.-V., F.B. and A.A. undertook laboratory work for the experiments and materials described in this study.

P.C. and P.K. established the analysis methods for the axonal transport experiments.

B.N., T.M.C., N.C.-R. and R.W.-M. conceived the study and experimental design and contributed to data analysis and interpretation. T.M.C., N.C.-R. and R.W.-M. supervised the study. B.N. drafted the manuscript. B.N., M.I.S., D.P.O., N.B.V., S.A.C. and R.W.-M edited the manuscript. R.W.-M. finalised the manuscript.

## Acknowledgements

We thank Dr Tina Wei, Prof Zameel Cader and Prof John B. Davis for sharing the MEA equipment used in this study; Prof Dominic Walsh for sharing a vial of rabbit serum containing the S97 anti-Aβ antibodies; the Oxford Brain Bank for facilitating the request for frozen brain tissues; Dr Emma Whiteley and Prof Colin Akerman for sharing their cortical neuron differentiation protocol; Drs Roman Fischer and Iolanda Vendrell from the Oxford Discovery Proteomics Facility for their help with MS data acquisition. Lastly, we would like to thank all donors of brain tissues and cell lines for this study.

---

**Supplementary Table 1:**
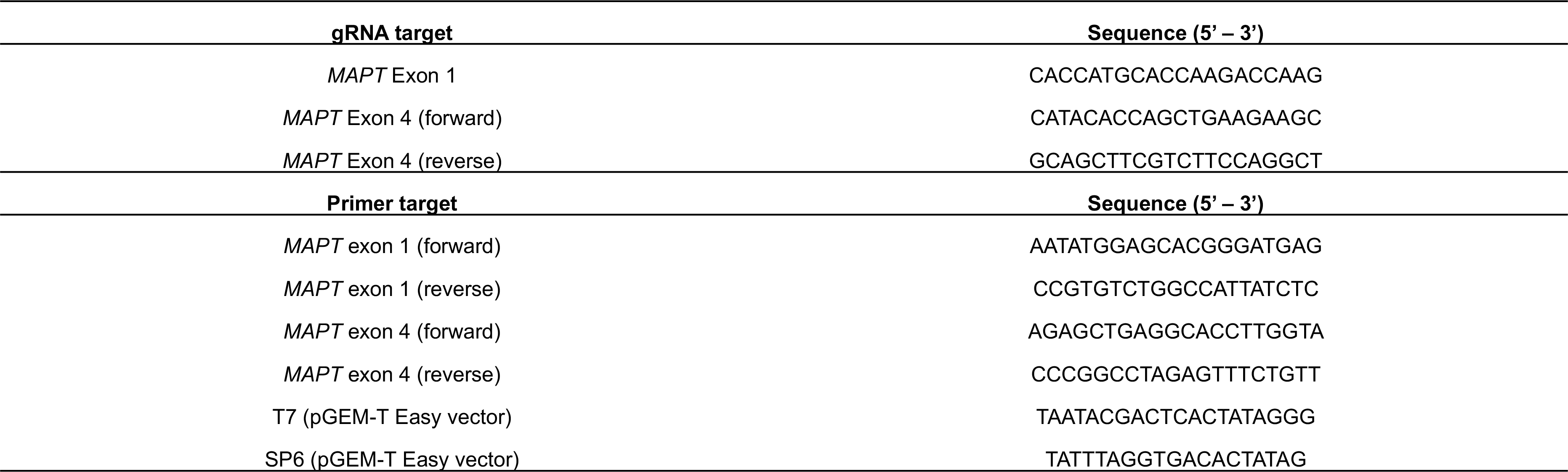
gRNA used (shown as DNA sequence) for targeting the *MAPT* gene with CRISPR-Cas9 system and sequencing primers spanning the targeted regions.

**Supplementary Table 2:** Primary and secondary antibodies used for Western blot and immunocytochemistry.

| **Target of primary antibodies for Western blot** | **Dilution factor** | **Manufacturer and catalogue number** |
| --- | --- | --- |
| Tau-1 | 1:1000 | Merck MAB3420 |
| Tau-5 | 1:500 | Abcam ab80579 |
| Tau-46 | 1:500 | Santa Cruz sc-32274 |
| Actin conjugated to HRP | 1:20,000 | Abcam ab49900 |
| **Secondary antibody for Western blot** | **Dilution factor** | **Manufacturer and catalogue number** |
| Goat anti-mouse IgG (H+L) HRP conjugate | 1:5,000 | Bio-Rad 1706516 |
| **Target of primary antibodies for immunocytochemistry** | **Dilution factor** | **Manufacturer and catalogue number** |
| Synapsin I/II | 1:500 | Synaptic Systems 106004 |
| Homer1 | 1:500 | Synaptic Systems 160003 |
| MAP2 | 1:1,000 | Abcam ab92434 |
| Human nuclear antigen | 1:200 | Abcam ab191181 |
| Cleaved caspase 3 | 1:500 | Abcam ab2302 |
| CUX2 | 1:200 | Abcam ab130395 |
| TBR1 | 1:200 | Abcam ab31940 |
| CTIP2 | 1:500 | Abcam ab18465 |
| **Secondary antibodies for immunocytochemistry** | **Dilution factor** | **Manufacturer and catalogue number** |
| Goat anti-guinea pig Dylight 488 | 1:1,000 | Abcam ab96959 |
| Goat anti-mouse Alexa Fluor 488 | 1:1,000 | Thermo A11001 |
| Goat anti-rabbit Alexa Fluor 555 | 1:1,000 | Thermo A27039 |
| Goat anti-chicken Alexa Fluor 555 | 1:1,000 | Thermo A21437 |
| Goat anti-rabbit Alexa Fluor 647 | 1:1,000 | Thermo A21245 |
| Goat anti-rat Alexa Fluor 647 | 1:1,000 | Thermo A21247 |

**Supplementary Table 3:**
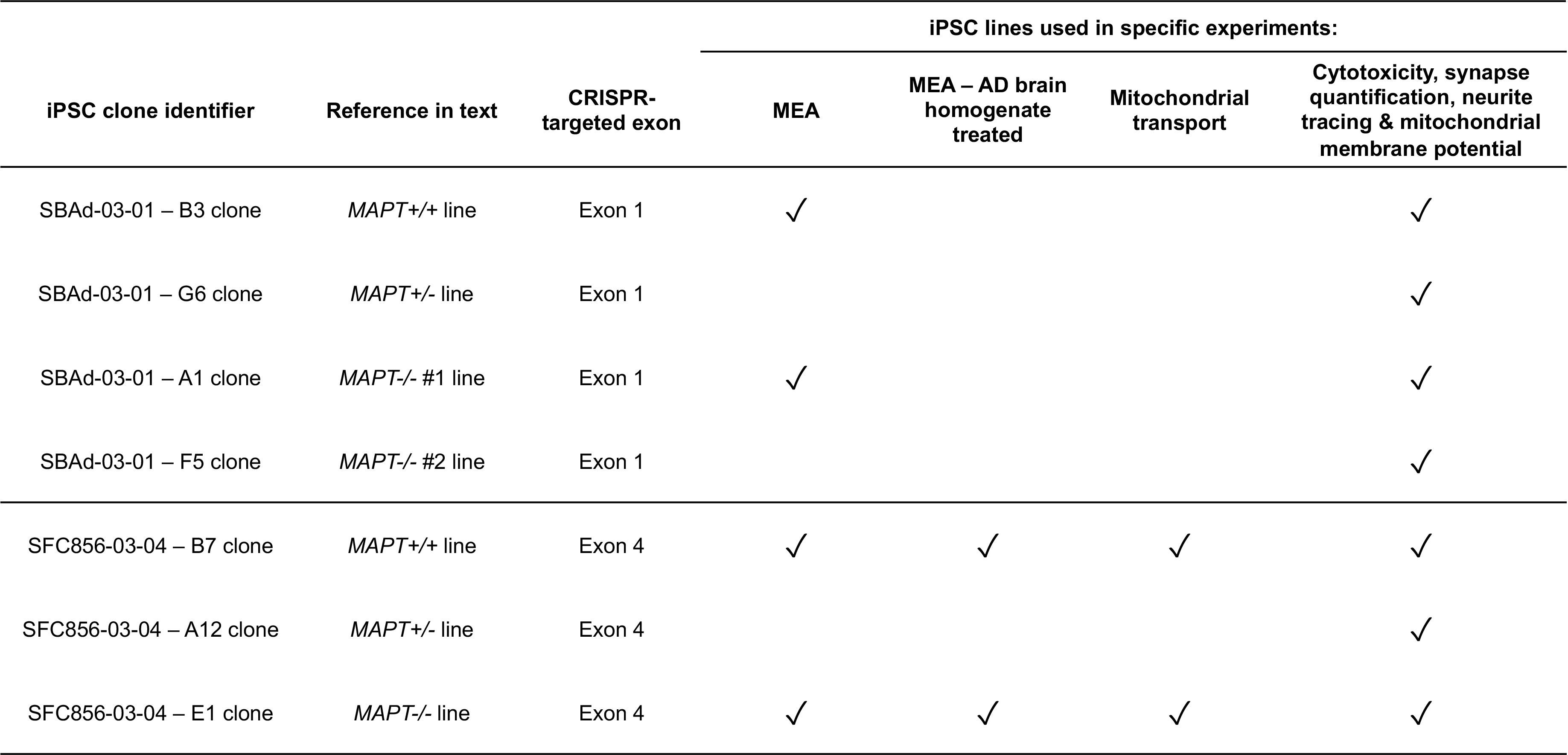
Specific iPSC lines used for downstream experiments on iPSC-derived cortical neurons.

| iPSC clone identifier | Reference in text | CRISPR-targeted exon | iPSC lines used in specific experiments: |  |  |  |
| --- | --- | --- | --- | --- | --- | --- |
|  |  |  | MEA | MEA – AD brain homogenate treated | Mitochondrial transport | Cytotoxicity, synapse quantification, neurite tracing & mitochondrial membrane potential |
| SBAd-03-01 – B3 clone | *MAPT*<sup>+/+</sup> line | Exon 1 | ✓ |  |  | ✓ |
| SBAd-03-01 – G6 clone | *MAPT*<sup>+/-</sup> line | Exon 1 |  |  |  | ✓ |
| SBAd-03-01 – A1 clone | *MAPT*<sup>-/-</sup> #1 line | Exon 1 | ✓ |  |  | ✓ |
| SBAd-03-01 – F5 clone | *MAPT*<sup>-/-</sup> #2 line | Exon 1 |  |  |  | ✓ |
| SFC856-03-04 – B7 clone | *MAPT*<sup>+/+</sup> line | Exon 4 | ✓ | ✓ | ✓ | ✓ |
| SFC856-03-04 – A12 clone | *MAPT*<sup>+/-</sup> line | Exon 4 |  |  |  | ✓ |
| SFC856-03-04 – E1 clone | *MAPT*<sup>-/-</sup> line | Exon 4 | ✓ | ✓ | ✓ | ✓ |

**Supplementary Figure 1:**
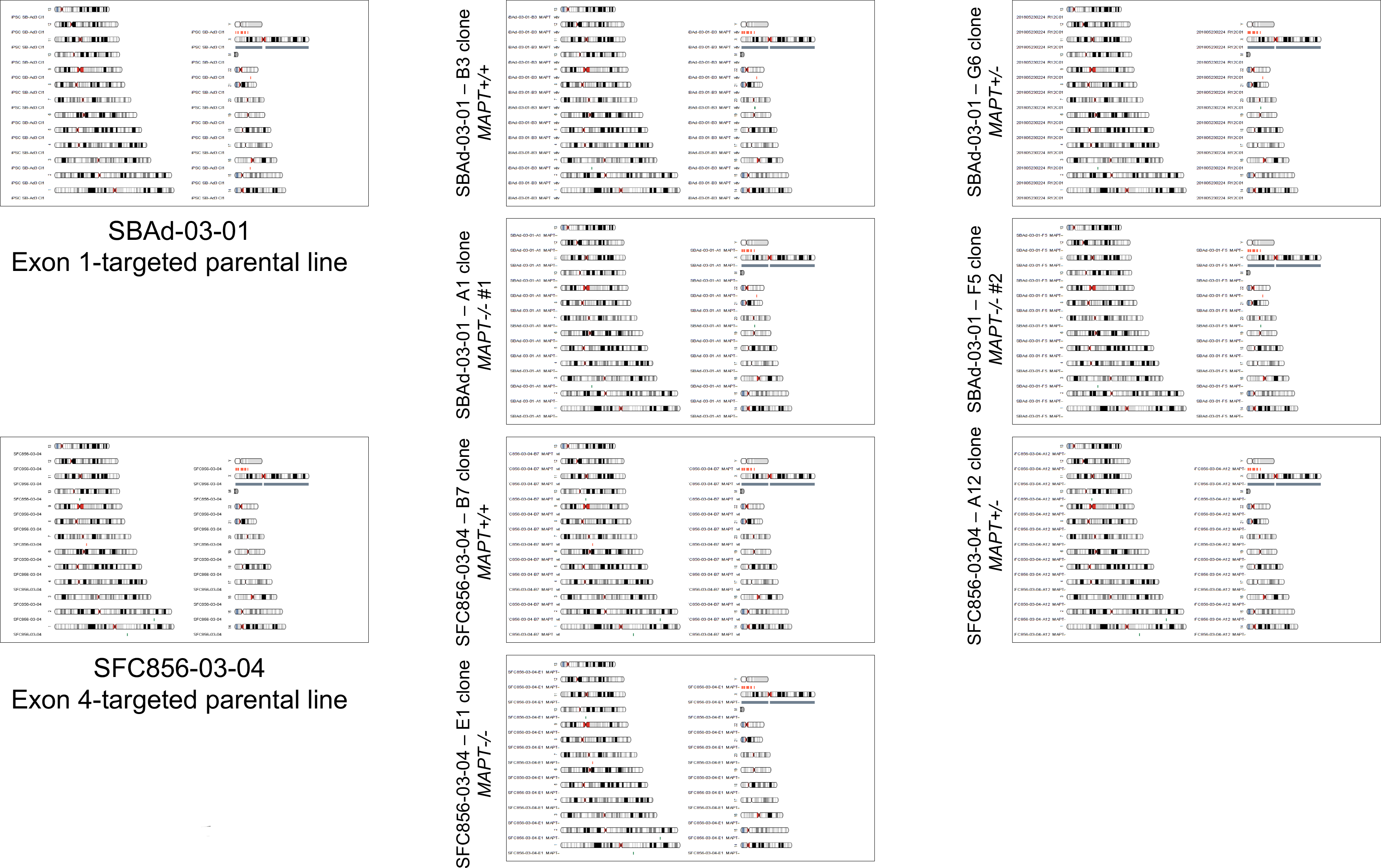
Quality control of iPSC lines used in this study. Genome integrity of the parental iPSC lines and both isogenic panels that underwent CRISPR-Cas9 gene editing targeting *MAPT*, examined by the Illumina OmniExpress24 single nucleotide polymorphism array (except the parental SBAd-03-01 line which was examined by the Illumina CytoSNP-12 array). Karyograms (KaryoStudio, Illumina) show amplifications (green)/deletions (orange)/loss of heterozygosity regions (grey) alongside the relevant chromosome. Female X chromosome is annotated in grey.

**Supplementary Figure 2:**
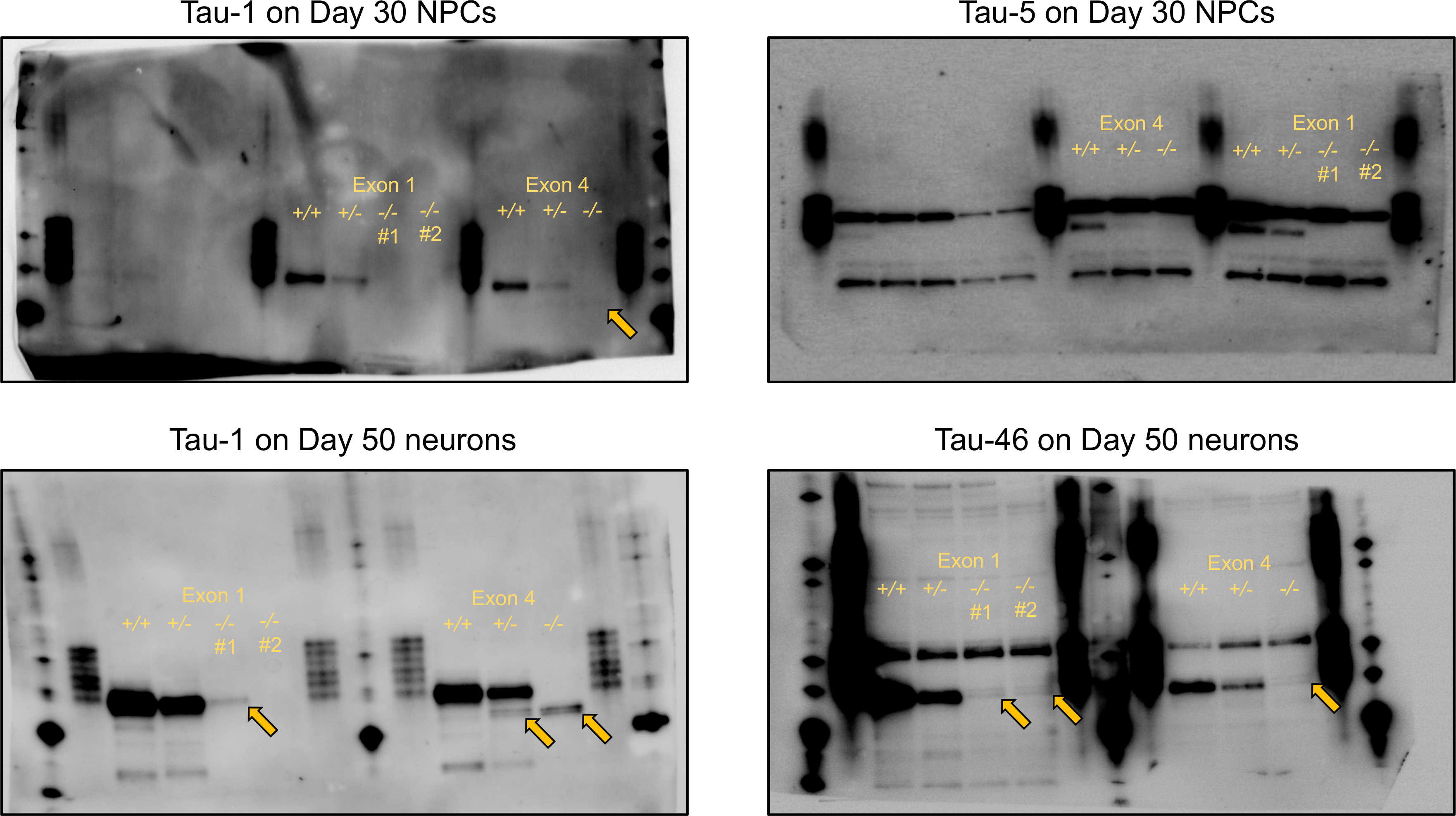
Full Western blots as in Fig. 1D with artificial signal augmentation. Western blots with either Day 30 NPCs or Day 50 neurons for all seven iPSC lines used in this study probed by Tau-1 (mid-region), Tau-5 (mid-region) and Tau-46 (C-terminus) antibodies. The bands in question are indicated by yellow arrows. Recombinant tau ladders flanked the labelled lines.

**Supplementary Figure 3:**
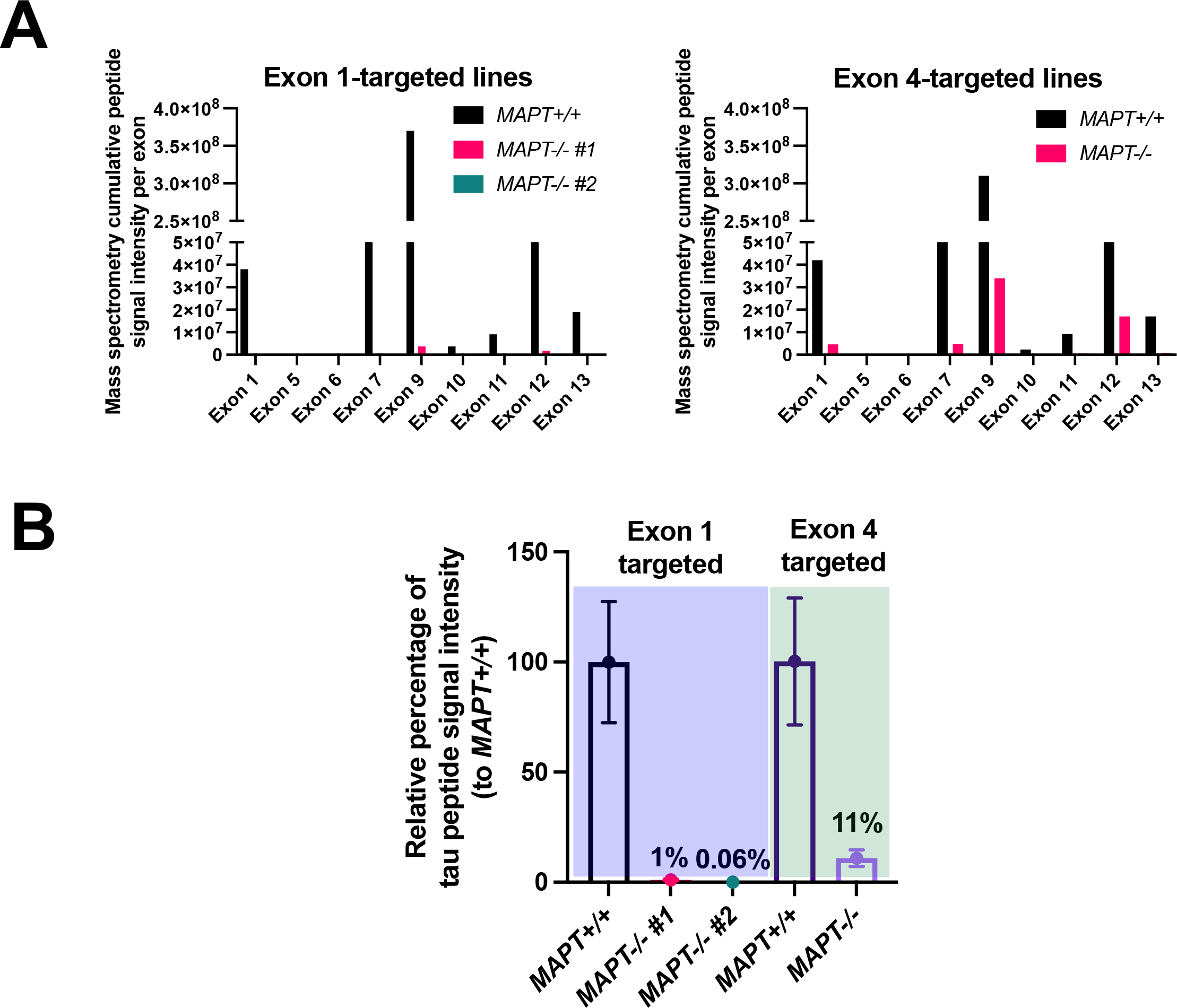
IP-MS experiment identified tau peptides present in Day 50 *MAPT-/-*neurons. **(A)** Quantifications of cumulative tau peptide signal intensity per exon identified from the IP using polyclonal K9JA antibody (Dako) binding to the C-terminus of tau in the *MAPT+/+* neurons compared to the *MAPT-/-* neurons. **(B)** Quantifications of relative tau peptide signal intensity percentage to those in the respective *MAPT+/+* neurons. Mean ± SEM. *n* = 51 tau peptides identified in the *MAPT+/+* neurons from the IP-MS analysis.

**Supplementary Figure 4:**
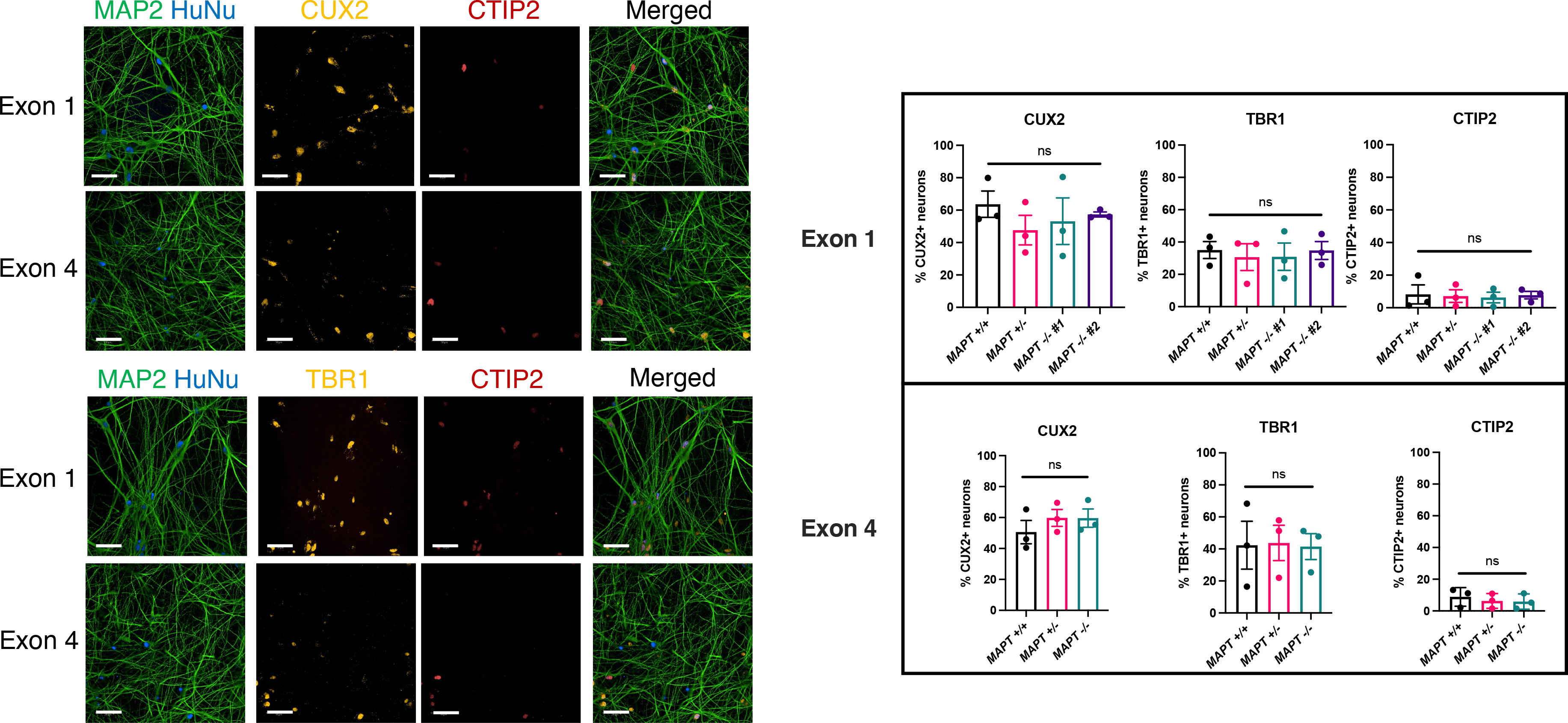
Cortical marker expression levels of the iPSC-derived neurons used in this study. Representative fluorescent images of Day 80 iPSC-derived cortical neurons from the *MAPT+/+* lines in co-culture with primary rat cortical astrocytes (Scale bar = 50 μm), as well as quantifications showing the quantification of cortical marker expression levels in Day 73-80 (Exon 1) or Day 80-84 (Exon 4) neurons. CUX2 (upper cortical layer), TBR1 (deeper cortical layer) and CTIP2 (deeper cortical layer, early marker) were used for staining in human nuclear antigen-positive cells. Mean ± SEM. *n* = three independent cortical neuron differentiation repeats. Kruskal-Wallis test with Dunn’s multiple comparison correction was used for statistical analysis.

**Supplementary Figure 5:**
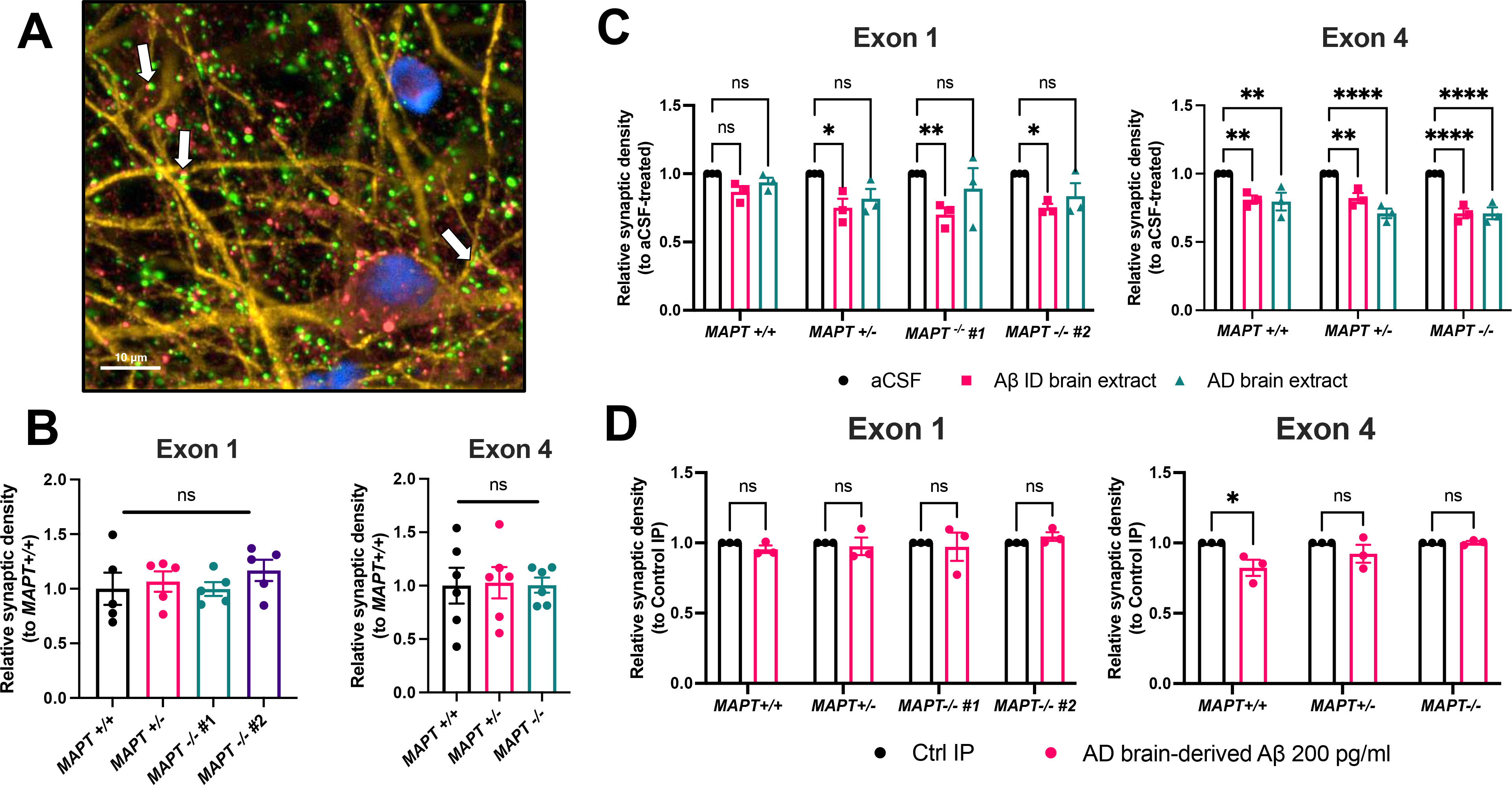
Aβ-driven synapse loss in iPSC-derived cortical neurons. **(A)** Representative fluorescent image of Day 88 iPSC-derived cortical neurons labelled with presynaptic Synapsin I/II (green), postsynaptic Homer1 (red), MAP2 (yellow) and human nucleus (blue). Synapsin I/II and Homer1 spots were identified in apposition within MAP2-positive regions to be considered a synapse indicated by white arrows. Scale bar = 10 μm. **(B)** Quantifications of relative synaptic density in Day 74-89 iPSC-derived cortical neurons compared to the respective *MAPT+/+* line <u>at baseline</u>. Mean ± SEM. *n* = five (Exon 1) or six (Exon 4) independent neuronal differentiation repeats. Kruskal-Wallis test with Dunn’s multiple comparison correction was used for statistical analysis. **(C)** Quantifications of relative synaptic density in Day 74-89 iPSC-derived cortical neurons <u>treated with 12.5% (v/v) AD or Aβ-ID brain homogenate for five days compared to the control aCSF treatment within each genotype. Mean ± SEM. *n* = three independent neuronal differentiation repeats. Two-way ANOVA with Dunnett’s multiple comparison correction was used for statistical analysis.</u> **(D)** Quantifications of relative synaptic density in Day 79-86 iPSC-derived cortical neurons <u>treated with either 200 pg/ml of AD brain-derived Aβ obtained from IP using antibodies targeting Aβ</u> or equal volume of eluate from control IP using normal mouse IgG for five days. Mean ± SEM. *n* = three independent neuronal differentiation repeats. Two-way ANOVA with Šídák’s multiple comparison correction was used for statistical analysis.

**Supplementary Figure 6:**
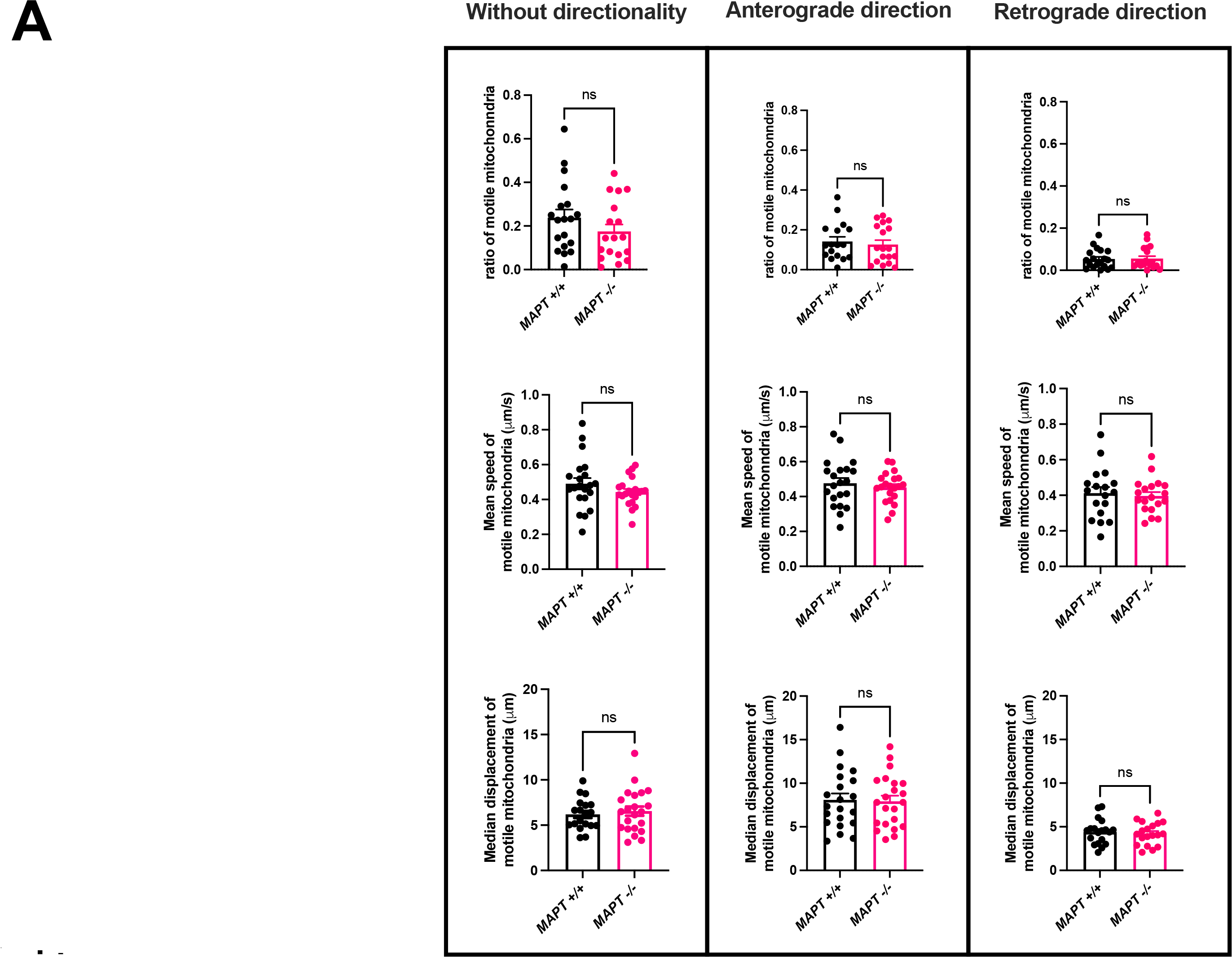
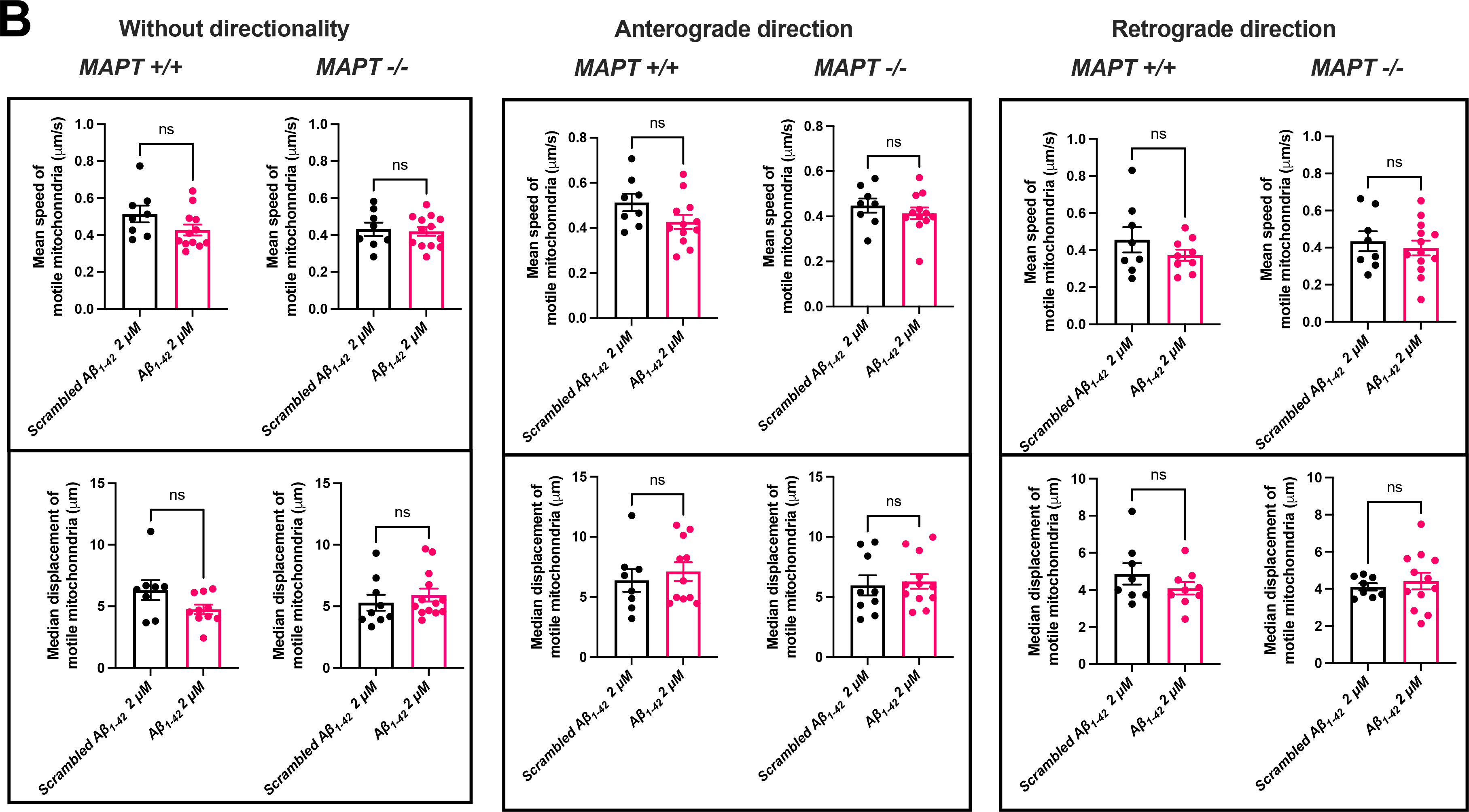
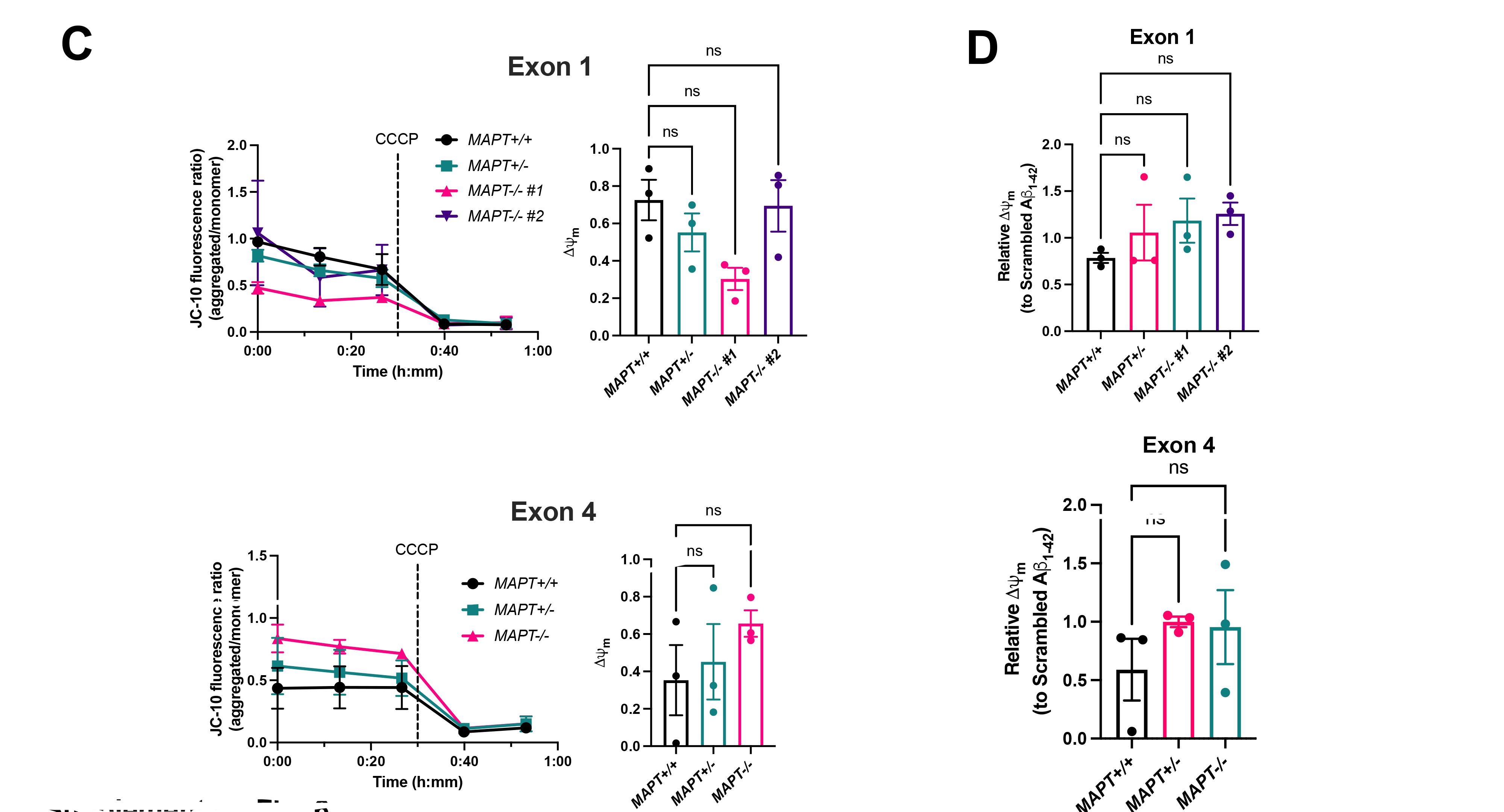
Axonal transport of mitochondria and their membrane potential in iPSC-derived cortical neurons. **(A)** Quantifications of ratios (to stationary mitochondria), speed and displacement of motile mitochondria <u>at baseline</u> with or without directionality over 150 s of live imaging in Day 70-95 iPSC-derived cortical neurons from the Exon 4 isogenic panel. Mean ± SEM. *n* = 17-21 (*MAPT+/+*) or 18-22 (*MAPT-/-*) microfluidic chambers measured across five independent neuronal differentiation repeats. Two-tailed Mann-Whitney test was used for statistical analysis. **(B)** Quantifications of speed and displacement of motile mitochondria at treated with either 2 μM scrambled Aβ<u>_1-42_</u> or Aβ<u>_1-42_ oligomers for 1 h</u> before imaging with or without directionality over 150 s of live imaging in Day 70-95 iPSC-derived cortical neurons from the Exon 4 isogenic panel. Mean ± SEM. *n* = 8 (*MAPT+/+* scrambled Aβ_1-42_), 13 (*MAPT+/+* Aβ_1-42_), 6-9 (*MAPT-/-* scrambled Aβ_1-42_) and 12-14 (*MAPT-/-* Aβ_1-42_) microfluidic chambers measured across five independent neuronal differentiation repeats. Two-tailed Mann-Whitney test was used for statistical analysis. **(C)** Quantifications of relative mitochondrial membrane potential (ψm) measured with JC-10 dye in Day 52-56 iPSC-derived cortical neurons from both isogenic panels <u>at baseline</u> compared to the *MAPT+/+* line. Mean ± SEM. *n* = three independent neuronal differentiation repeats. Kruskal-Wallis test with Dunn’s multiple comparison correction was used for statistical analysis. **(D)** Quantifications of relative mitochondrial membrane potential (ψm) measured with JC-10 dye in Day 52-56 iPSC-derived cortical neurons from both isogenic panels <u>treated with either 2 μM scrambled Aβ</u>_1-42_ <u>or Aβ</u>_1-42_ <u>oligomers for 1 h</u> before imaging compared to the *MAPT+/+* line. Mean ± SEM. *n* = three independent neuronal differentiation repeats. Kruskal-Wallis test with Dunn’s multiple comparison correction was used for statistical analysis.

**Supplementary Figure 7:**
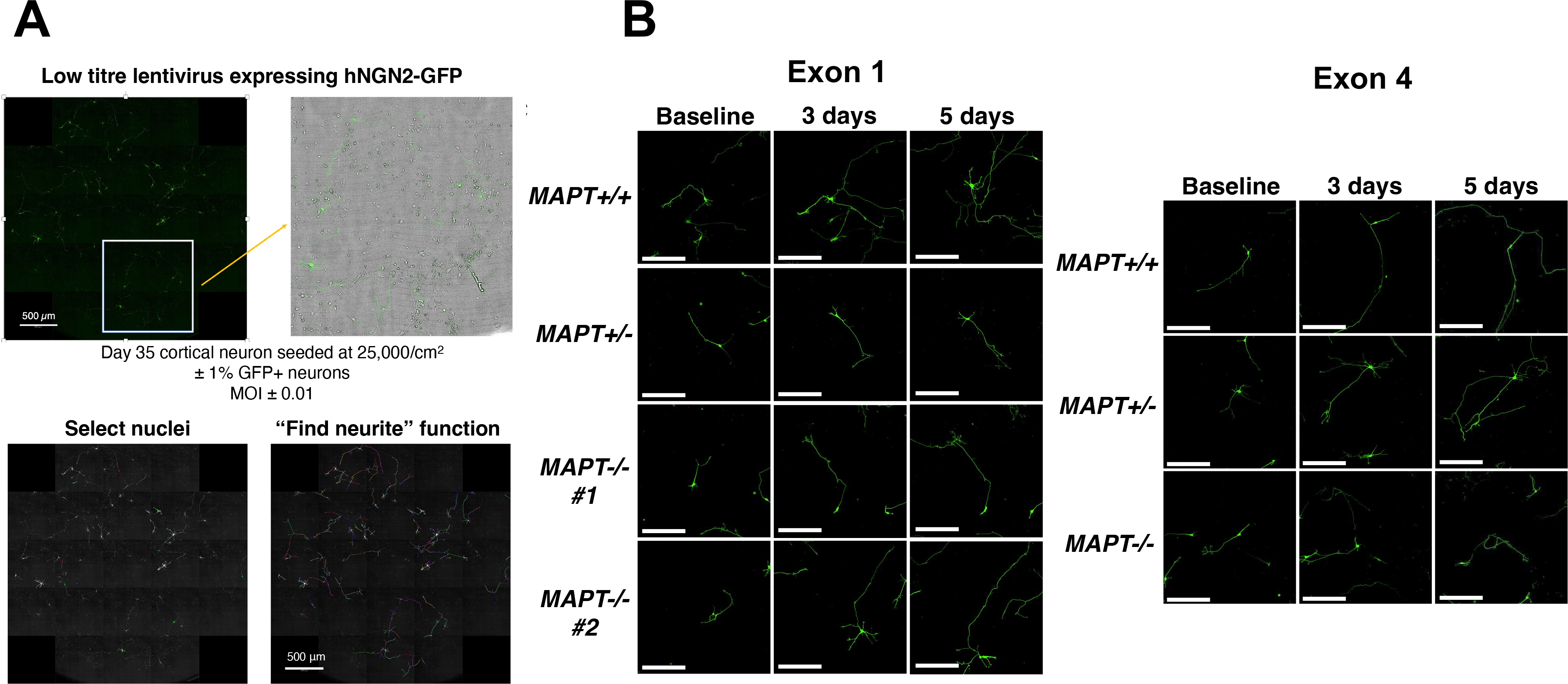
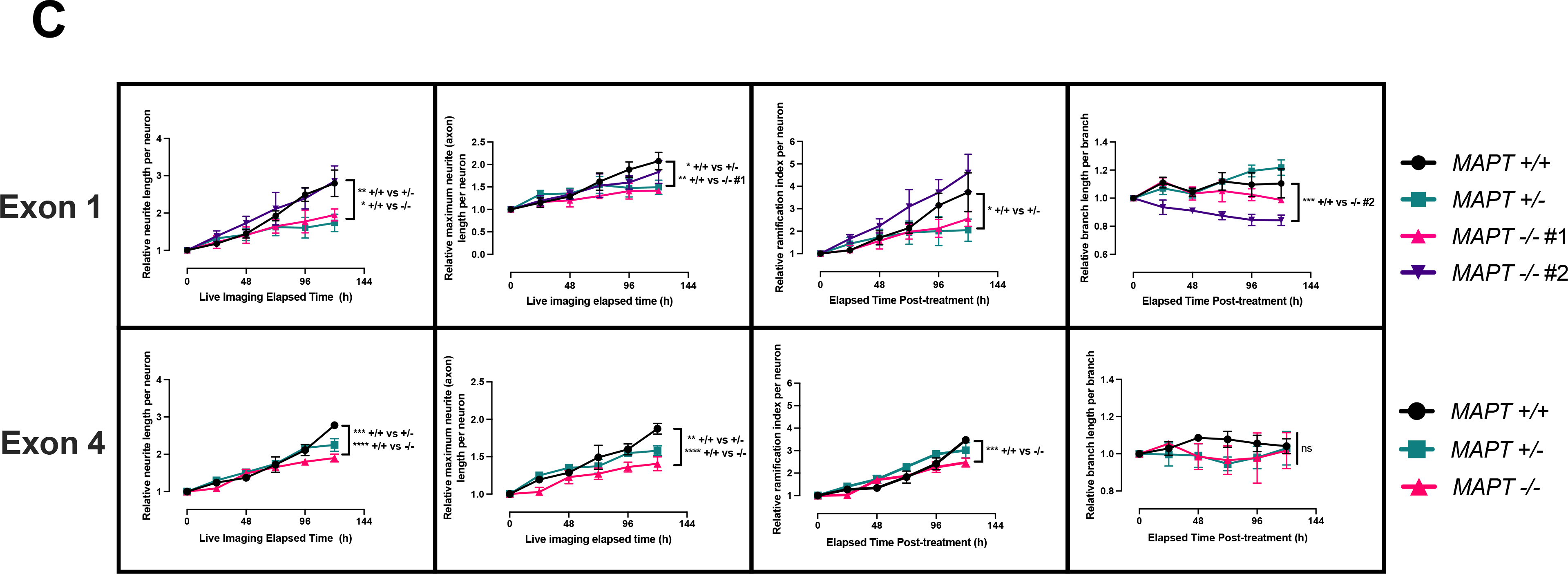
Neurite outgrowth impairment in *MAPT-/-* neurons except the Exon 1 *MAPT-/-* #2 line. **(A)** Schematic of neurite tracing experiment design. The images show a subset of Day 35 iPSC-derived cortical neurons expressing GFP, and in the Harmony^®^ software analysis pipeline identifying GFP-positive nuclei and neurites stemming from those nuclei. Scale bar = 500 μm. **(B)** Representative fluorescent images of Day 35 iPSC-derived cortical neurons in both isogenic panels demonstrating neurite outgrowth over 5 days of live imaging. Scale bar = 100 μm. **(C)** Quantifications of relative neurite outgrowth, axonal outgrowth, ramification index and neurite branch length in Day 35 iPSC-derived cortical neurons over 5 days of live imaging compared to the baseline (i.e., elapse time = 0 h). Mean ± SEM. *n* = three independent neuronal differentiation repeats. Two-way ANOVA with Dunnett’s multiple comparison correction was used for statistical analysis.

